# Optogenetic therapy: High spatiotemporal resolution and pattern recognition compatible with vision restoration in non-human primates

**DOI:** 10.1101/2020.05.17.100230

**Authors:** Gregory Gauvain, Himanshu Akolkar, Antoine Chaffiol, Fabrice Arcizet, Mina A. Khoei, Mélissa Desrosiers, Céline Jaillard, Romain Caplette, Olivier Marre, Stephane Bertin, Claire-Maelle Fovet, Joanna Demilly, Valérie Fradot, Elena Brazhnikova, Philippe Hantraye, Pierre Pouget, Anne Douar, Didier Pruneau, Joël Chavas, José-Alain Sahel, Deniz Dalkara, Jens Duebel, Ryad Benosman, Serge Picaud

## Abstract

Restoring vision using optogenetics is an ideal medical application because the eye offers a direct window to access and stimulate the pathological area: the retina. Optogenetic therapy could be applied to diseases with photoreceptor degeneration such as retinitis pigmentosa. Here, we select the specific optogenetic construct that is now used in the clinical trial and assess the opsin functional efficacy on non-human primate’s retinal ganglion cells (RGCs).

We chose the microbial opsin ChrimsonR and showed that the vector AAV2.7m8 produced greater transfection in RGCs compared to AAV2, and that ChrimsonR attached to tdTomato (ChR-tdT) is more efficiently expressed than ChrimsonR. The 600 nm light activates the RGCs transfected with the vector AAV2.7m8-ChR-tdT from an irradiance of 10^15^ photons.cm^-2^.s^-1^. Vector doses of 5.10^10^ and 5.10^11^ vg/eye transfect up to 7000 RGCs/mm^2^ in the perifovea, with no significant immune reaction. Furthermore, using a multielectrode array we recorded RGCs responses starting from 1ms stimulus duration. Using the recorded activity we were able to decode stimulus information and estimate a theoretical visual acuity of 20/249, above legal blindness. Altogether, our results pave the way for the ongoing clinical trial with the AAV2.7m8-ChrimsonR-tdT vector for vision restoration in patients affected by retinitis pigmentosa.

**One Sentence Summary:** We select here the vector and genetic construct best suited to provide vision restoration in patients suffering from retinopathies, we demonstrate temporal resolution compatible with high dynamic visual scenes and a visual acuity above legal blindness.

## Introduction

Optogenetics has transformed neurobiology by giving scientists the ability to control the activity of excitable cells using light^1^. Optogenetic therapy has also created great hope for new forms of brain-machine interface with cell selectivity and distant optical control. Rebuilding vision though optogenetic approaches is conceptually straightforward as it aims at reintroducing light sensitivity in the residual retinal tissue after photoreceptor degeneration in diseases like retinal dystrophies^2^ or age-related macular degeneration^3^. Since these diseases mainly affect photoreceptors, the remaining retinal layers are still available to communicate with the brain through the optic nerve. Retinal prostheses have already proven the feasibility of reactivating these retinal layers^4,5^ despite their major limitations in surgery, spatial resolution and cell specificity^6^.

The use of optogenetic to restore vision was first proposed by Zhao Pan. His team expressed the microbial opsin Channelrhodopsin-2 (Chr2) in RGCs of blind mice^7^ and subsequently in the retina of normal marmosets^8^. These studies led to a clinical trial using this microbial opsin but no result has been published since its start in February 2016^9^. Other retinal cell types upstream in the circuit were subsequently targeted to restore vision in blind rodents, postmortem retinal tissue and non-human primates: bipolar cells^10–12^ and dormant cone photoreceptors^13,14^. Clinically, the choice of the targeted cell type will depend on the stage of tissue remodeling following photoreceptor degeneration^15–17^. In a translational effort, we decided to focus our efforts on targeting retinal ganglion cells (RGCs) - the neurons projecting their axons out of the retina - because this strategy could work for all patients having lost their photoreceptors, regardless of the disease stage^18^.

RGC activation can be obtained in the non-human primate retina using the more sensitive form of Chr2, -‘CatCH’^19^ This was tested under a RGC specific promoter^20^. However, the required blue light intensities were close to radiation safety limits^21^. It was thus clinically important to evaluate other potentials opsins conferring a balance between light sensitivity and channel kinetics^22^.

Here, we selected the optimum AAV capsid with the most red-shifted opsin, with a wavelength 45 nm longer than ReaChR^23^, and we demonstrate that its high spatiotemporal resolution is suitable for visual restoration. A single intravitreal injection at 5.10^10^ or 5.10^11^ vg/eye transfects up to 7000 RGCs/mm^2^ in the perifovea. Responses were elicited for 1ms stimulus duration and saturated at 30-50 ms stimulus durations. Furthermore, responses generated on a multielectrode array enabled us to identify moving bars and letters with a theoretical visual acuity of 20/249, which is higher than the threshold for legal blindness. These characterizations of visual response in the non-human primate retina pave the way for the ongoing clinical trial with the AAV2.7m8-ChrimsonR-tdT vector for vision restoration in patients affected by retinitis pigmentosa.

## Results

### Subhead 1: AAV2.7m8-ChR-dtT provides the greatest efficacy of transduction

Our first objective was to determine the best genetic construct for expressing ChrimsonR in primate RGCs. It is known that intravitreal delivery of AAV vector in non-human primates. (NHPs) leads to the transduction of the ganglion cell layer in the peri-foveal ring^20,24^. Because the mutated capsid AAV2.7m8 has shown a stronger transduction of the peri-fovea^25^, we decided to compare its efficacy in expressing ChrimsonR (ChR) with that of the wild type AAV2 version. Then, as ChR is often used fused to the fluorescent protein tdTomato to visualize its cellular expression, we also assessed whether the native protein ChR and the fused protein Chrimson-tdTomato were equally expressed in primate RGCs. The four selected constructs (AAV2 and AAV2.7m8 vectors encoding either ChR or ChR-tdT) were each injected in 4 different eyes at the same concentration (5×10^11^ vg/eye) in 8 animals (Table S1). The expression level of the microbial opsin was functionally assessed 2 months after the in-vivo intravitreal injection. The transduced retinas were isolated ex-vivo and divided in hemi-fovea for either large scale 256 multielectrode array (256-MEA) extracellular recordings or for 2-photon targeted patch-clamp recordings (Fig. 1). All recordings were done in the presence of synaptic blockers to suppress any natural light responses (see material methods). 256-MEA recordings revealed a great disparity in the efficacy of the different vectors to generate a functional expression of ChR (Fig. 1A-D). The quality of recording was defined by the number of electrodes where a spontaneous spiking activity could be measured (active electrodes: 152 +/− 46 electrodes per retina explants on average) while the ChR efficacy was defined by the number of electrodes with an increase in activity during presentation of light flashes (responsive electrodes, SN ratio >4). This quantification showed that there was a significant difference between the different constructs; with AAV2.7m8-ChR-tdT presenting the greatest efficacy (Fig. 1C, 64.4% of active sites are responsive vs 13.4%, 10.6% and 0% for AAV2.7m8 – ChR-tdT, AAV2.7m8 – ChR, AAV2 – ChR-tdT and AAV2 – ChR, respectively, *P*<0.001). For the construction with no tdT fluorescence, we could not confirm the location of ChR expression on the 256-MEA using tdT fluorescence. Therefore, the tissue was repositioned multiple times on the array to increase the sampling area when no light response was measured. Light sensitivity was measured using a range of light intensities, from 1.37×10^14^ to 6.78×10^16^ photon s.cm^2^.s^-1^ on all responsive retinas (Fig. 1B, Fig. 1D). AAV2.7m8 – ChR-tdT displayed responses in the 4 tested retinas (versus 2, 1 and 0 for AAV2.7m8 – ChR, AAV2 – ChR-tdT and AAV2 – ChR, respectively). It also showed the highest sensitivity with responses recorded for 2.34×10^15^ photons.cm^2^.s^-1^, and higher recorded frequency than the other construct. The increase in spiking activity as a function of light irradiance was observed from light onset and continued for the whole stimulation duration, which was characteristic of optogenetic activation. Furthermore an increase in the number of responsive electrodes was clearly visible while increasing the irradiance (Fig. 1B). We recorded the action spectrum of the responses (Fig. 1E), and the measured peak was in agreement with the known spectral sensitivity of ChR, around 575nm^23^.

**Fig 1:**
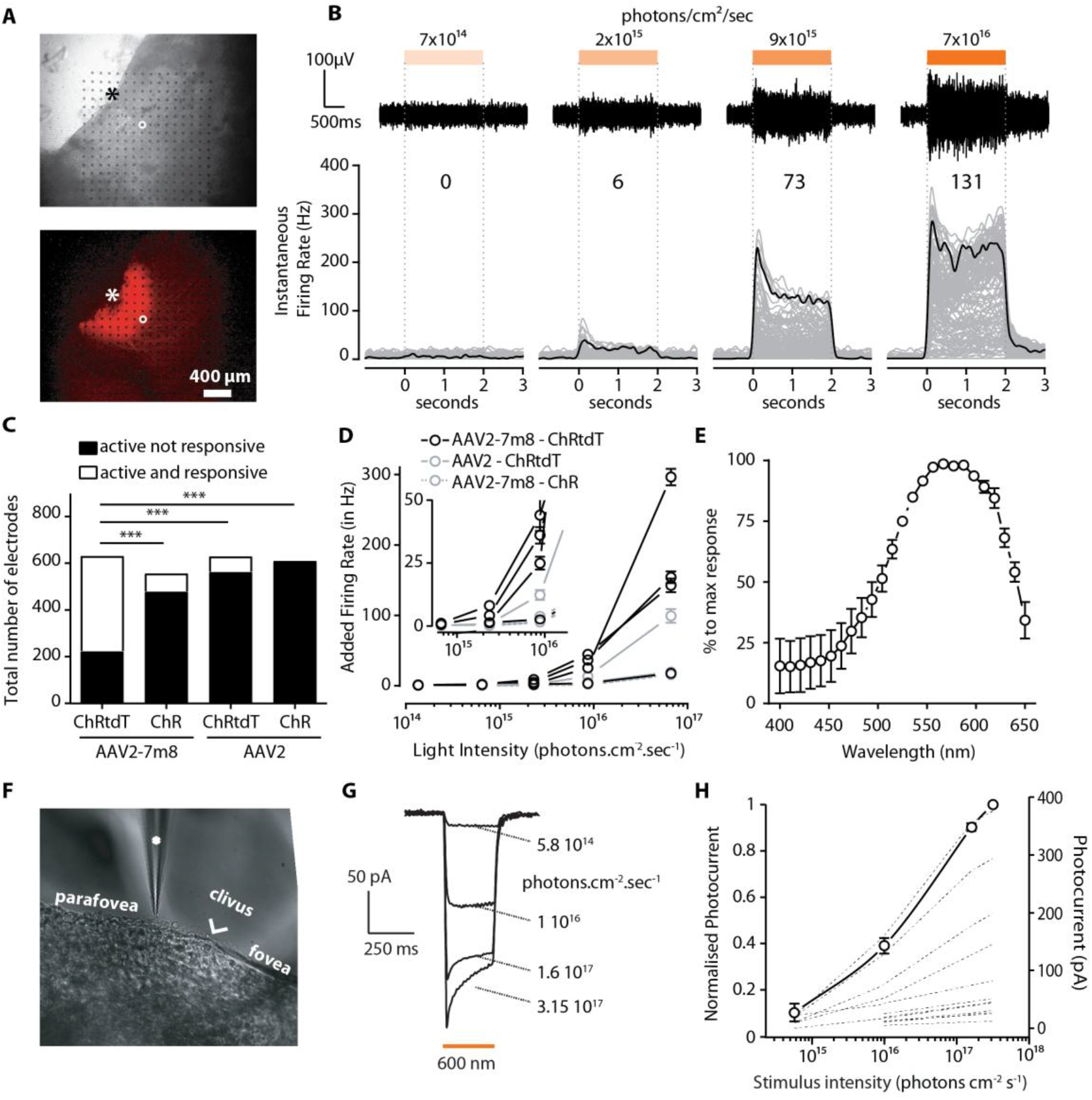
AAV2.7m8-ChRtdT demonstrate greater infectability in NHP retinas. **(A *top*)** Infrared image of a hemi-fovea from a primate injected with AAV2.7m8 – ChRtdT, as observed during MEA recording. Electrodes can be seen as black dots. *(**Bottom**)* Epifluorescence image of the same piece of retina. Strong peri-foveal expression can be observed. An asterisk denotes the fovea center. (**B *top***) Raw signal for one sample electrode (circled electrode in (A)) in response to stimuli of increasing intensities. *(**Bottom**)* Instantaneous firing rate as a function of time, before, during and after a 2 seconds stimulus for all electrodes of one hemi-fovea (grey lines, 197 lines. Firing rate is averaged for 10 repetitions. Black lines show the average firing rate for the electrode displayed in the top panels. The number on top indicates the number of responsive electrodes per light intensity. (**C**) Number of active electrodes summed for the 4 experiments in each different recordings conditions (theoretical maximum: 1024). Fraction of active and responsive electrode is maximal for AAV2-7m8 ChRtdT (*n* = 4, Fisher contingency test, *P*<0.001). (**D**) Average added firing per responsive retina for the four construct +/− SEM (no response for the AAV2-ChR construct). Stimulation at 590nm +/−15nm. Zoomed inset emphases first responsive intensity: 2.34 × 10^15^ photons.cm^-^2.s^-1^. (**E**) Average normalized action spectrum for three retina expressing AAV2.7m8 ChRtdT +/−SEM. (**F**) Infrared image of perifoveal region from a primate injected with AAV2-7m8 – ChRtdT recorded using 2 photon targeted patchclamp. The patch-clamp electrode is marked with a white asterisk. (**G** and **H**) Whole-cell patchclamp recordings of ChRtdT –expressing macaque peri-fovea neurons. (**G**) Photocurrent traces from one recorded cell at different light intensities. (**H**) ChR-induced photocurrents are represented as a function of light intensity (for each cell (n=17), dashed lines, and averaged after normalization, solid line). Light stimulation intensity ranged from 5.8 × 10^14^ to 3.2 × 10^17^ photons.cm^-2^.s^-1^.

Results obtained with 256-MEA experiments were confirmed through two-photon targeted patch-clamp recordings (Fig. 1F-H) using the other half of the foveae. AAV2.7m8 – ChR-tdT elicited, at the highest irradiance, robust responses with a typical photocurrent shape consisting of a fast transient followed by a steady-state current (Fig. 1G, 12 to 375 pA, average: 88.7 +/− 25.5 pA, *n*=17). These currents increased steadily when increasing the light intensity from 5.8×10^14^ up to 3.15×10^17^ photons.cm2.s^-1^ (Fig. 1H). With the AAV2.7m8 – ChR-tdT combination, we recorded 18 responsive cells (5, 0, 7, 6 cells/retina) whereas only 4 (0, 3, 1, 0 cells/retina) were obtained with the AAV2 – ChR-tdT construct (data not shown) and none for the others. For the without tdT fluorescence, AAV2.7m8 – ChR and AAV2 – ChR, recordings were performed on random healthy RGCs in the peri-foveal area (>40 cells per conditions). In these later conditions, none of the recorded RGCs showed light-evoked responses even under conditions known to activate ChR. Taken together, these results confirmed the superiority of the AAV2.7m8 – ChR-tdT construct, which was systematically used in the following sections.

### Subhead 2: AAV2.7m8 – ChR-tdT at 5×10^11^ provides greater light sensitivity at 6 months post-injection

Once the capsid and genetic payload selected we assessed the transgene stability in time with long term expression (6 months). In the same set of experiments we optimized virus load using three distinct vector quantities for our intravitreal delivery: 5 × 10^9^, 5 × 10^10^ and 5 × 10^11^ vector genome per eye (vg/eye), for a total of six animals (four eyes per dose). Results from 4 retina treated with the high dose were subsequently added. After the injections, we examined the animal’s eyes each month for posterior uveitis and vitreal haze. Clinical grade evaluation showed no significant immune response following ChR-tdT expression (fig. S1). The success of our vision restoration strategy relies on: 1) a large and dense area of transfected cells and 2) a high light sensitivity per cell. To correctly estimate the cell number and retinal coverage of the expression, we performed manual cell count in RGC layers on the confocal stack of images on hemi-foveas. Using these counts we established density maps (Fig. 2A) and density profiles (Fig. 2B). We observed an increased number of ChR-tdT expressing cells with increasing vector dose (averaged total transfected cells: 491 +/− 64,4395 +/− 631 and 5935 +/− 715, for ChR-tdT at 5 × 10^9^, 5 × 10^10^ and 5 × 10^11^ vg/eye, respectively, see methods). While the local density reached with the two highest concentrations does not differ statistically (Fig. 2A), eyes injected with 5 × 10^11^ vg express ChR-tdT towards moderately higher eccentricity (Fig. 2B), giving this dose a potentially larger area covered with expression. Using automated counting of DAPI stained nuclei in the same samples, we estimated the peak density to be ~40000 RGCs/mm2, at an eccentricity of 0.4mm (fig. S1), in agreement with previous RGC density maps^26^. This number allows us to estimate the fraction of RGCs expressing ChR-tdT to ~20%. These hemi-foveas were used, prior to fixation, for MEA recordings to test light sensitivity following longterm expression (Fig. 2C). In terms of fraction of responsive electrodes we see no clear differences between 5 × 10^10^ and 5 × 10^11^ vg/eye, but a reduced number for the lower dose (only 1 on 4 retina with responsive electrodes, Fig. 2D). More importantly, the different virus doses lead to different level of light sensitivity, with 5 × 10^11^ vg/eye having the strongest overall responses and lowest response threshold (Fig. 2E, see Table S2 for Tukey’s multiple comparisons test). Comparing 2 and 6 months expression time, we do not observe any differences in the fraction of responsive electrodes (Fig. 2D, 102 +/− 58 vs. 73 +/− 65 for 2 months and 6 months 5 × 10^11^ vg/eye, respectively), nor in light sensitivity (Fig. 1D and Fig. 2E). Due to this, and the absence of any significant immune response from the viral load or the ectopic gene expression (fig. S2), we selected 5 × 10^11^ vg/eye as the desired dose for our therapy. As such all subsequent data presented here are from 5 × 10^11^ vg/eye dose at 6 months time expression.

**Fig 2:**
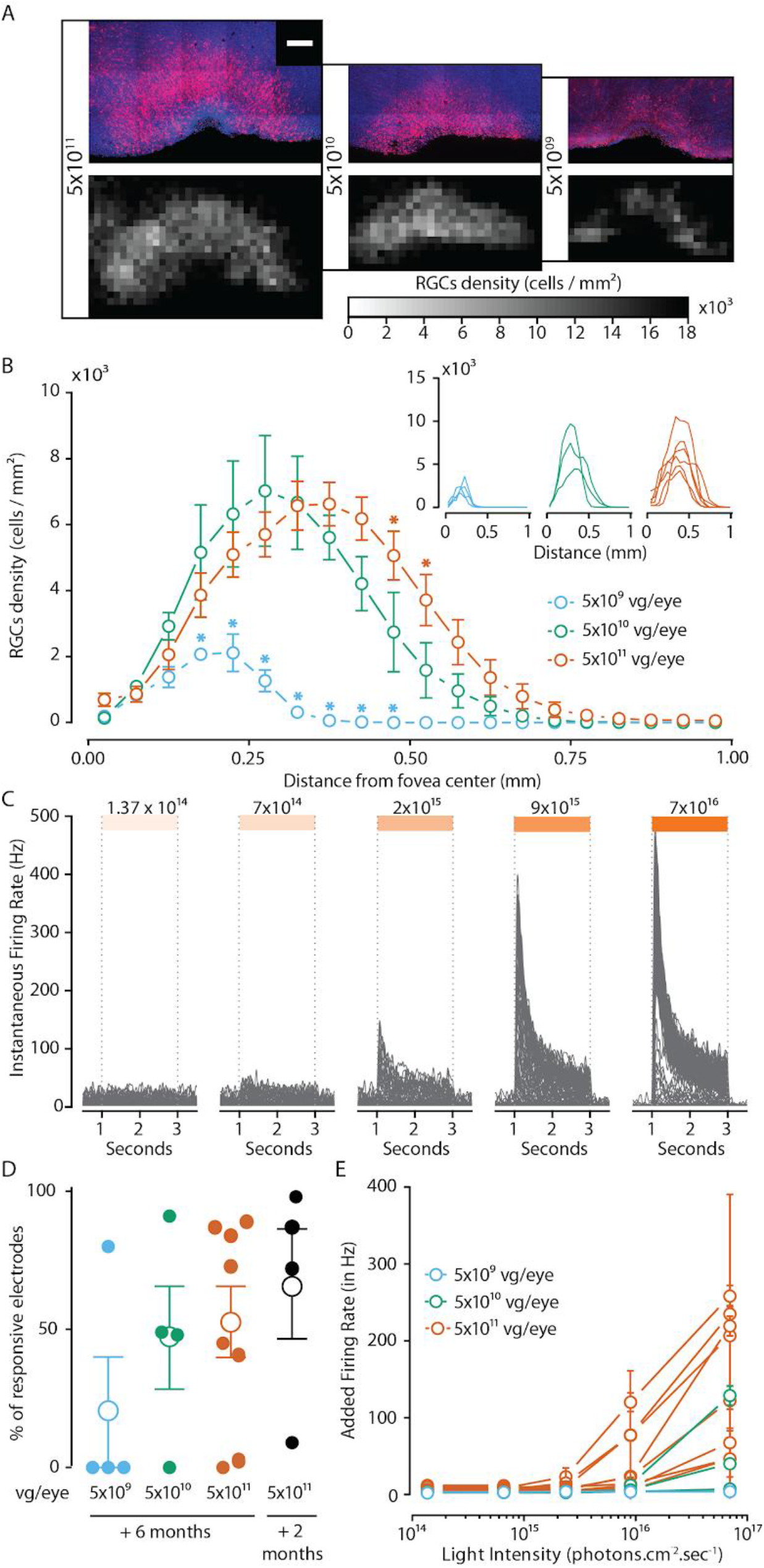
AAV2.7m8 – ChRtdT induced high density long-term expression in primate perifovea. **(A *left*)** Projections of confocal stack stitches for three different doses of vector, red: ChRtdT expressing cells, blue: DAPI nuclei staining. Scale bar: 200μm. *(**Right***) Density map of ChRtdT positive RGCs cells for the same three example hemi-foveas. (**B**) ChRtdT expressing RGCs density relative to retina eccentricity for the three vector dose tested for long term expression. Density is expressed as the average for all counted retinas (*P*<0.05, 2 way ANOVA multiple comparisons). Individual retina profile can be observed in the inset. **(C)** Spike density function for all responsive electrodes of one retina in response to different light levels (**D**) Fraction of responsive electrodes measured in MEA experiments for all conditions, all doses 6 months and 2 months expression at 5 × 10^11^ vg.eye (no significative differences, Wilcoxon-Mann-Whitney test). Plain circle: value for individual retina, open circle: average for all the retina +/− SEM. (**E**) Average added firing per responsive retina for the three doses and at different light level +/− SEM at 590nm +/−15nm, see Table S2 for dose response statistical analyses.

### Subhead 3: Activity modulation at the millisecond-scale

Natural vision relies on a high temporal dynamic range of information to perceive moving objects. In virtual reality goggles, the minimum information transfer mode appears to rely on video rate (30Hz, *27*). Therefore, vision restoration for locomotion or perception of dynamic scenes should bring back light sensitivity at least in this temporal scale. We therefore measured the temporal dynamic of our optogenetic responses using full-field monochromatic stimuli (2 × 10^17^ photons.cm^-2^.s^-1^ at 600 nm +/− 10 nm) of increasing duration (1 ms to 2000 ms). This light intensity was selected because it generated highest firing rates while remaining below radiation safety limits for continuous eye exposure (~6 × 10^17^ photons.cm^2^.s^-1^^28,29^). Significant light responses were detected for stimulus durations starting at just few milliseconds (Fig. 3A). Interestingly the firing rate of RGCs reached a plateau for durations ranging between 30 to 100 ms depending on the tested retina (Fig. 3B). To define the minimal stimulus duration generating a reliable response, we computed time to first spike following stimulation onset for all responsive electrodes (Fig. 3C). From the 5 ms stimulus and thereafter, we observed a median time to first spike around 9 ms. This indicates that 5ms of stimulation is enough to activate RGCs reliably, and that intracellular integration of ChR-tdT photocurrent initiated spiking in less than 10 ms for most of the responsive electrodes. Furthermore, for stimulation duration of 20 ms time to first spike for 50% of the responsive electrodes is comprised between 3 and 11 ms. We then looked at the distribution of firing rates following stimulation of increasing duration (Fig. 3D). Even for 1 ms stimuli (Fig. 3C-D, dark blue curves), 12% of electrodes measured a peak firing rates exceeding 100Hz. Even though we were considering here multi-unit recording, this observation indicates that, for some RGCs, 1 ms stimulus was sufficient to elicit a strong response as clearly seen in Fig. 3A. For 5 and 20 ms stimuli, 48% and 69% of the overall responsive electrodes had firing rates above 100Hz, respectively. Finally, for the longest tested stimulus duration (2s), peak responses decreased and the sustained firing rate was also reduced during consecutive stimulations (Fig. 3A,B, D) but both recovered subsequently. To further investigate this effect for long stimulation duration, we computed the Fano factor, a measure of the variability in the number of spikes in relation to the mean number of spikes, for all electrodes as a function of stimulation duration. While the Fano factor is below 1 for the shorter duration, indicative of lower variability than Poisson distribution, we observed a dramatic increase in the spike train variability for 2 seconds stimulations (Fig. 3E). Most of this effect can be attributed to the stimuli hysteresis as the retinal sensitivity was subsequently recovered. However, it suggests that limiting optogenetic stimulation to temporally discrete stimuli is recommended to generate strong and reliable cell responses. These stimuli should be composed of flickering stimuli to reduce the total amount of light and risk of increased spike train variability. For a light pulse width between 5 and 20 ms, the frequency of stimulation could be comprised between 100 and 25 Hz.

**Fig 3:**
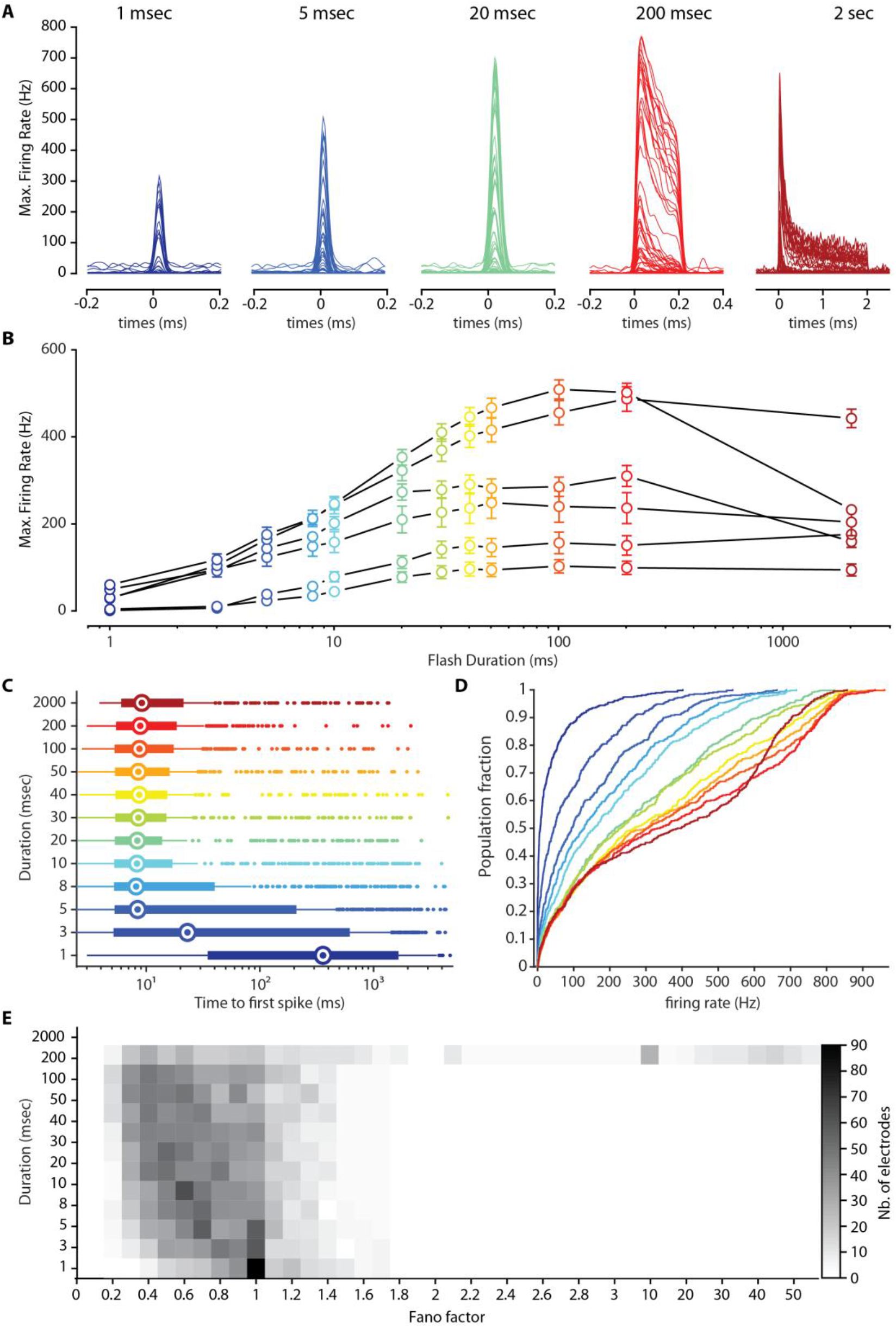
millisecond activation of ChR-tdT expressing primates RGCs. (**A**) Spike density function for all responsive electrode of one retina in response to increasing duration (1 – 5 – 20 – 200 ms and 2s, left from right). (**B**) Average maximal firing rate measured for retinas treated with 5 × 10^11^ vg/eye for stimuli of increasing duration and constant light intensity (n=6, 2 × 10^17^ photons.cm^-2^.s^-1^, 600nm +/−10nm). (**C**) Vertical box plot displaying time to first spike following stimulation onset as a function of stimulus duration. Recordings from the different retinas are pooled, as such each electrode as the same weight. Medians are displayed as a hollow circle, box edges are the 25^th^ and 75^th^ percentiles, whiskers extend to the extreme points, outliers plotted individually. (**D**) Cumulative plot of maximal firing rate per electrode versus stimulus duration, duration color coded as in (**C**). (**E**) Fano factor as a function of stimulation duration for all responsive electrode-Value of 1 represent poisson distribution, value below 1 indicate an increase in information content (**C, D** and **E**: *n=488* electrodes from 6 retinas expressing ChR-tdT).

### Subhead 4: ChR-tdT can produce high temporal photocurrent and spiking modulation

In parallel to our population study on MEA, we examined the temporal dynamic at the single cell level by recording photocurrent modulation at the single cell level. In all recorded hemi-fovea, fluorescent transfected cells were discernible in the characteristics half torus shape (Fig. 4A-B). Using two-photon guided patch-clamp techniques, we recorded ChR-tdT expressing RGCs in the peri-fovea using cell-attached or voltage-clamp intracellular configuration (Fig. 4C). We first replicated the analysis of photocurrent modulation by light intensities to compare responses at 6 months (Fig. 4C to E) and 2 months of expression (see Fig. 1F-H). We observed that averaged and normalized photocurrent followed a similar photo-sensitivity curve (Fig. 4D), with activation threshold in the 10^15^ photons.cm^-2^.s^-1^ intensity range, and robust responses to light stimuli at a wavelength of 600 nm (+/− 10 nm) well below the illumination radiation safety limits for the human eye^21,29,30^. When comparing peak photocurrent or peak firing rate for maximal light intensity (3.15 × 10^17^ photons.cm^-2^.s^-1^), we found no statically significant difference between the two expression durations (Fig. 4E), suggesting a stable expression of ChR-tdT in this timeframe.

**Fig 4:**
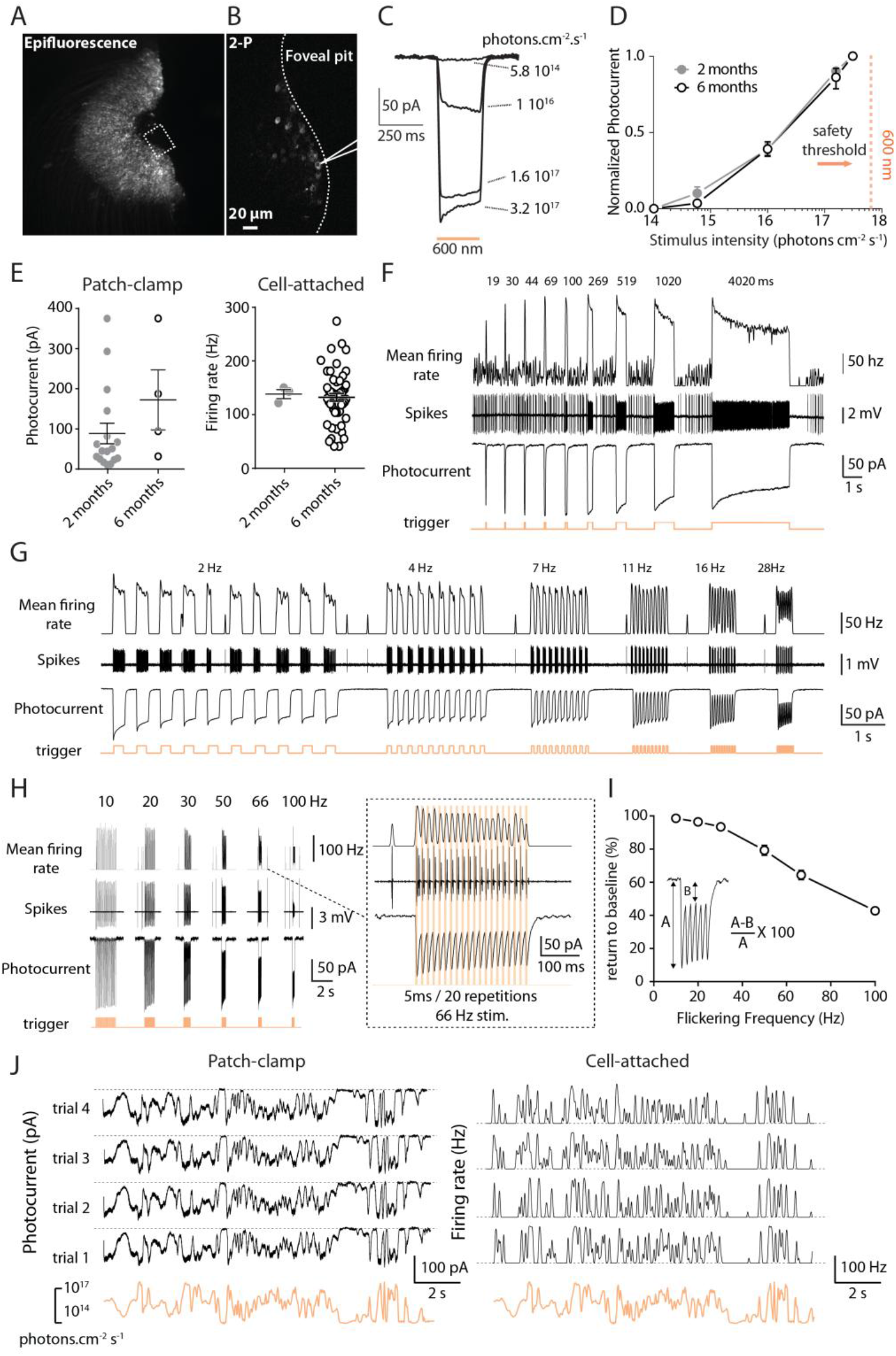
long-term stable expression of ChRtdT allows reliable spike train generation at high temporal resolution. (**A**) Epifluorescence image showing tdT expressing RGCs in the peri-foveal region of a monkey 6 months after injection at 5 × 10^11^ vg/eye. (**B**) Two-photon laser imaging of the foveal pit, scanned zone is presented as a dotted rectangle in A, patch-clamp electrode outline visible in vicinity of a ChRtdT-positive cell. (**C**) Photocurrent responses of a representative cell to different light intensities (5.8 × 10 to 3.2 × 10 photons.cm^-2^.s^-1^), obtained by patch-clamp recording. (**D**) Comparison of normalized photocurrent response versus irradiance at 600nm +/− 10nm in ChRtdT-expressing RGCs following a 5 × 10^11^ vg/eye (*n*=17 for 2 month, *n*=4 for 6 month). (**E**) Peak photocurrent (***left***) and maximum firing rate (***right***) in ChR-tdT expressing RGCs for the two durations, 2 and 6 months (photocurrent: 88.7 +/− 25.5 pA, *n*=17 and 172.4 +/− 74.9 pA, *n*=4, respectively, p=0.155; firing rate: 138.3 +/− 8.4 Hz, *n*=3 and 132.3 +/− 7.2 Hz, *n*=50, respectively, p=0.669). (**F *Top***) Firing frequency in response to stimuli of increasing flash duration obtained in cell attached configuration; average of two repetitions. (**F *middle***) Raw data showing the spiking behavior of the recorded cell. (**F *bottom***) Photocurrents recorded in the same cell during presentation of the same pattern of stimulations in whole cell configuration. The orange line at the very bottom of the figure shows the duration of the flash (19 ms to 4020 ms). (**G**) Firing frequency *(top),* spike trains (middle), and photocurrent response *(bottom)* of the same cell to increasing stimulus frequency in full duty cycle (from 2 to 30 Hz). (**H *left***) Firing frequency (*top*) and photocurrent response (bottom) of the same cell to increasing stimulus frequency, using short light stimulation pulses (trains consisting of 20 repetitions of a 5 ms stimulation), ranging from 10 to 100 Hz. (**H *right***) recorded RGC precisely follow a 66Hz stimulation by firing distinct action potentials doublets at every pulse. (**I**) Inactivation of the photocurrent as observed in (G) as a function of stimulation frequency. For every recorded cell (*n*=7) we measured the percentage of return to the current baseline (averaged across the 20 stimulations), from 2 to 100 Hz stimulations. For 10, 66 and 100 Hz stimulations, cells returned to 98.6 +/− 0.4 %, 64.4 +/− 2.9 %, and 42.8 +/− 1.8 % of their initial baseline value, respectively. Light intensity used in panel D to H is 3.2 × 10^17^ photons cm^-2^ s^-1^ at 600nm +/−10nm. (**J**) Representative example of photocurrent responses (left) and firing frequency (right) of the same cell, displaying the high reliability across trials (*n*=4) for both current and firing activity, while stimulating the cell with a randomized oscillating stimulus, intensities ranging from 3 × 10^14^ to 3 × 10^17^ photons cm^-2^ s^-1^.

To investigate the response kinetics, we recorded light responses to stimuli of increasing durations (Fig. 4F) in the cell attached mode (spikes) and in the whole cell configuration (photocurrent). The photocurrent and the spike rate both follow stimuli precisely and form a mirror image of one another. Interestingly, the reduced photocurrent amplitudes from its initial peak to smaller sustained amplitude are also mirrored by similar decrease in firing rates. We further investigated the effect of flicker stimuli either in 50% duty cycle (half of the stimulus period with the light ON, at 2 to 28 Hz) or at a specific stimulus duration (5 ms, from 10 to 100 Hz) (Fig. 4G-I). Using full duty cycle stimulation (Fig. 4G), the photocurrent closely followed the stimulus frequency up to a 30Hz flicker stimulation. These results are in agreement with the fast opening and closing kinetics of the ChR channel in RGCs (10 to 90% rise time, 5.2 +/− 1.7 ms; decay time, 27 +/− 2.9 ms for 250 ms duration stimuli at 3.15 × 10^17^ photons.cm^-2^.s^-1^1, *n*=5). The fast photocurrents allow neurons to translate robustly each light pulse into burst of spikes but the decay time of the photocurrent does not allow a complete return to the resting level during the stimulus trains (e.g. 30Hz flicker, Fig. 4G). We then used lower duty cycle, using trains of 5 ms stimuli (20 pulses at frequencies between 10 and 100 Hz) as it has been showed to largely activate ChR-tdT expressing RGCs in MEA experiments (Fig. 3). With such short stimuli our recordings showed that photocurrent could be modulated at high frequencies with large amplitudes (50-100 pA) and generate periodic spiking activities. Recorded cells followed precisely the stimulus train even at 100 Hz but current deactivation was incomplete between light pulses (Fig. 4H and I). Cell-attached recordings confirmed the ability of the neuron to follow the stimulus frequencies up to 66 Hz, even under conditions that elicited an important amount of incomplete current deactivation (Fig. 4I). This 60 Hz range would fit well with flicker perception limits as observed for natural vision in human subjects^31,32^ and can possibly be used for fast video rate stimulation in human patients. Finally, we activated cells using a stimulus with natural properties simulating a one dimensional random walk consisting in a rapidly changing contrast stimulus (full-field stimulus with intensities ranging from 3 × 10^14^ to 3 × 10^17^ photons cm^-2^ s^-1^). We observed a strikingly high response reliability across trials (*n*=4) for both current and firing rate activities (Figure 4-J). These results demonstrated that RGCs expressing ChR-tdT can follow a high dynamic range of visual stimulation compatible with human perception.

### Subhead 5: ChR-TdT can generate high spatial precision for visual perception

We have previously shown that the RGC responses followed precisely the temporal resolution of optogenetic stimuli. To test the spatial sensitivity of the optogenetic stimulation, the retina on the MEA was stimulated by circular spots of varying sizes (25um, 50um and 100um) centered on the MEA electrodes (100 um electrode pitch, 10 μm diameter) at a light intensity of 2.10 × 10^17^ photons.cm^-2^.s^-1^ (600nm +/− 10nm) (Fig. 5). Multi-unit electrode based analysis showed that even the electrodes that were far (up to 1mm) from the stimulated spot elicited an increase in spiking frequency (Fig. 5A), we performed spike sorting on the electrode signals to obtain single cell activity using unsupervised sorting algorithm. This spike sorting indicated that individual spikes were recorded on several electrodes as a consequence of spike propagation in RGCs axons running at the retinal surface toward the optic disk (Fig. 5A & fig. S2). The increase of latency with distance to the stimulated area was consistent with an anterograde propagation of spikes along axons. Taking advantage of this spike propagation, we measured the spike velocity in the ChR-tdT expressing cells (fig. S3). The unimodal distribution peaking at 0.5 m/s (fig. S3H) indicates that the ChR-tdT expressing RGC population contains a majority of midget RGCs^33^. This conclusion on the cell identity was consistent with the midget cell morphology of tdTomato expressing cells obtained by two-photon microscopy (Fig. S2A-E). However, very few cells (*n*=9) had faster velocities of axonal spike propagation (>1m/s) indicative of possible presence of parasol RGC in the ChR-tdT expressing RGCs.

**Fig 5:**
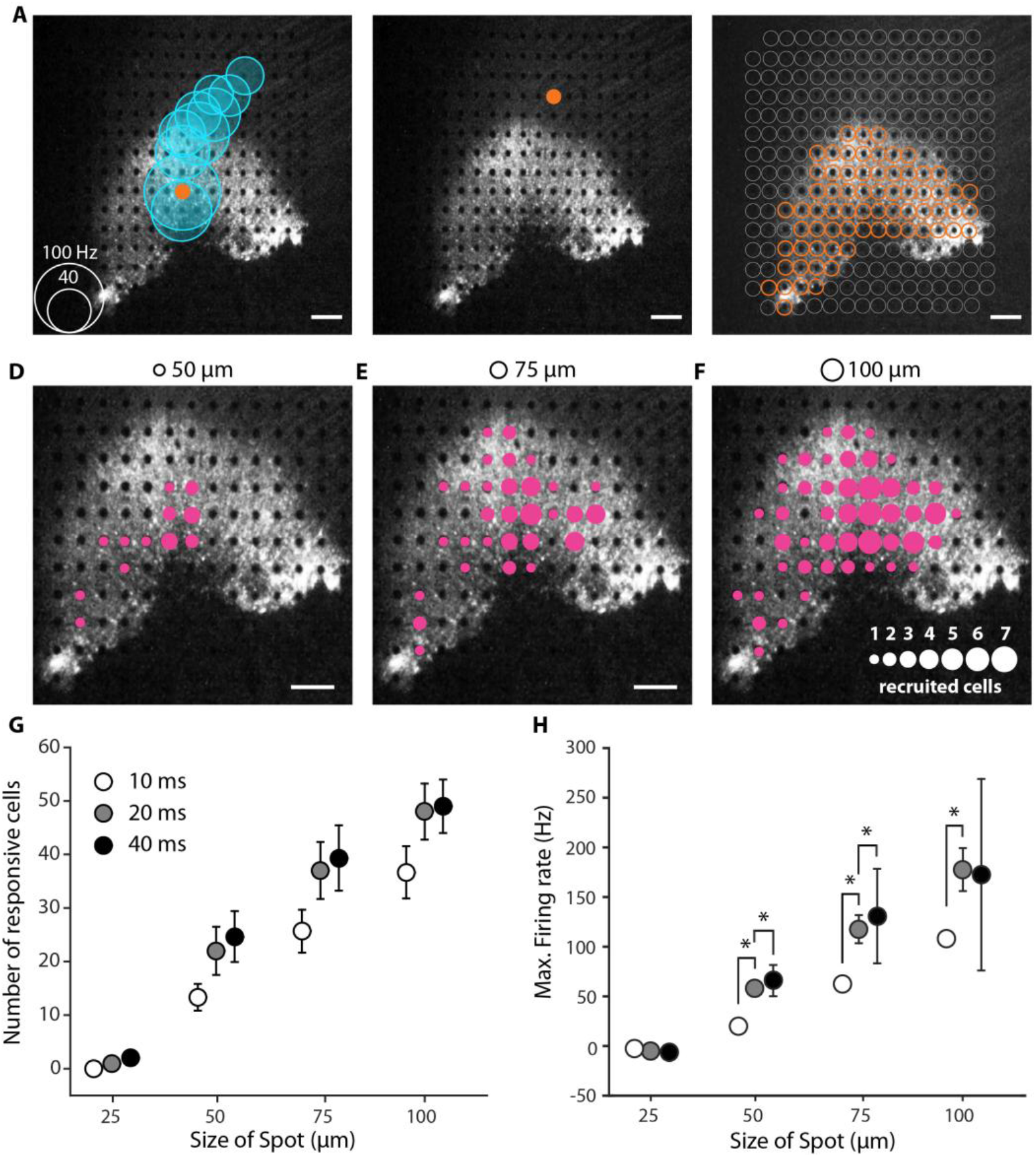
Spatio-temporal resolution for discrete stimulation. (**A**) Example of a spot stimulation 100 μm and 40 ms on the transfected peri-fovea showed strong propagating activity. Black and white image shows the fluorescence present in peri-fovea, electrodes can be seen as black dots. Orange dot shows the location of stimulation and the blue circles show the channels where spiking response was recorded. The size of the circle represents the maximum spiking rate inside a 100 ms window after stimulation, scale on bottom left corner. (**B**) Same as in A, except the stimulation is on the RGCs axons and not on the soma, no response measured. (**C**) All location where stimulation resulted in spiking response recorded on at least one MEA channel. These location matches exactly with the intense fluorescence in the perifovea. (**D-F**) Images showing the extent of responses for 20 ms presentation of spot stimuli at different sizes (displayed on the top of the images). Sizes of the magenta circles show the number of cells activated by a spot at that particular position see scale in the lower right part on panel (**F**). The total recorded cells (**G**) and mean firing rate (**H**) over the entire electrode array when size and spot duration were modulated. For all stimuli durations (10 ms, 20 ms, and 40 ms) the increase in size from 25 to 50 μm, 50 to 75 μm and 75 to 100μm led to statistically significant increase in the firing rate of the cells not shown on the figure (all p < 0.005), for duration increase at a single spot size results of statistical analyses are displayed on the figure. All scale bar: 200μm.

As tdTomato fluorescence was present in RGCs axons, we examined whether light stimulation could elicit spikes directly in ChR-tdT expressing axons, with anterograde and/or retrograde propagation. When a spot of light was centered on an electrode in contact with ChR-tdT positive axons but not ChR-tdT-expressing soma, we observed no increase in spike activity in any neighboring or distant electrode (Fig. 5B). This result demonstrated that optical stimulation of ChR-tdT expressed in axons was not sufficient to trigger spikes. In fact, a high correspondence was found between the area containing cell bodies expressing TdTomato and the location of electrodes with optogenetic responsive cells (Fig. 5C). When varying spot size and presentation duration we observed single cell activation for spots as small as 50 μm (Fig. 5D-F, Movie S1). The number of responsive cells and their spiking frequencies were dependent upon the spot size and the stimulus duration (Fig. 5G-H, fig. S4). It is worth noting here that our stimulations were centered on the opaque MEA electrodes (10um diameter), which could considerably reduce light intensity for the smaller spot (25μm in diameter). Nonetheless,

In order to assess the functional impact of our visual restoration strategy we tested the ability of treated retinas to encode information about direction and speed of motion, as well as the ability to discriminate patterns. First, we presented moving bars (75um wide) at various velocities (2.2 mm/s, 4.4 mm/s) with different orientations and directions across the treated retina (Movie S2). Based on the known retinal magnification factor, 1 arc degree of visual size corresponds to a size of 211 μm on the retina^34^, hence our bar stimuli correspond to a visual field angle of 0.375 degrees moving at 11 or 22 degrees/seconds. In order to compute the elicited visual flow, we used spike sorting on the recorded activity, then plane fitting on the peak of the cells responses to estimate the direction of the bar (Fig. 6, A and fig. S4) and speed (Fig. 6D). The plane fitting method allowed us to identify for each direction the unique succession of cell activation on the bar path (Fig. 6, A to C). This temporal response of the cells was found to be sufficient to correctly estimate the direction and velocity of the bar over the retina despite the discrete spacing of electrodes.

**Figure 6:**
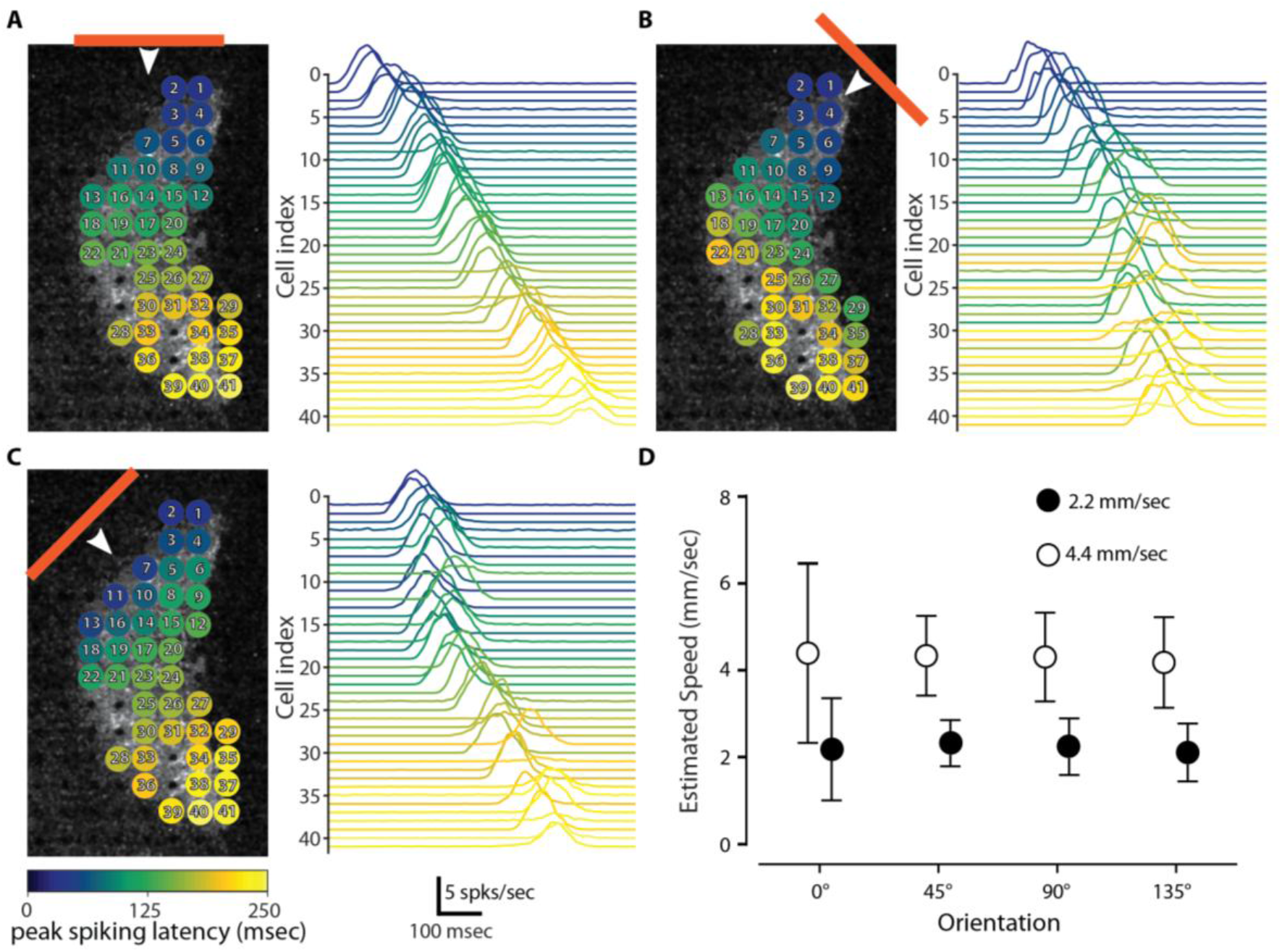
measured activity is enough to estimate speed and direction of a moving bar stimulation. Figure shows the distinct temporal patterns of the cells for a 75 um bar moving at 2.2 mm/sec across the retina at various orientations (**A, B, C**). The colors represent the relative time of peak spiking activity for each cell from the peak of earliest cell activation for the motion of the bar. The spike rate plots show the trial wise average spiking frequency response of the cells for each condition. (**D**) A simple plane fitting based method was able to predict the relative speed of the bars presented at either 2.2mm/sec or 4.4 mm/sec. Results averaged over 3 retinas samples.

The most widely used clinical tests to measure visual acuity, the Snellen chart, assess patient performance in reading letters, and can be used to evaluate vision restoration strategies^35^. In order to estimate an expected visual acuity our strategy could restore, we stimulated the retinas with different optotypes (X shape, circle and square) of various sizes from 55 to 330 μm (symbols width). All presented characters were moved on the retina explants through the fovea center (8 different directions, 50 trials each, randomized presentations, Movie S3). Following spike sorting of the recorded activity (Fig. 7A), we used an algorithm (^36^, see methods for details) to discriminate the tested letters of similar size directly from the generated spatiotemporal spike responses of the ChR-tdT expressing cells (Fig. 7B). Our results indicate an 83% discrimination rate for a symbol size of 220 μm (Fig. 7C, edge of symbol: 44 μm), in agreement we find maximal value for mutual information for 220μm (Fig. 7D).

**Figure 7:**
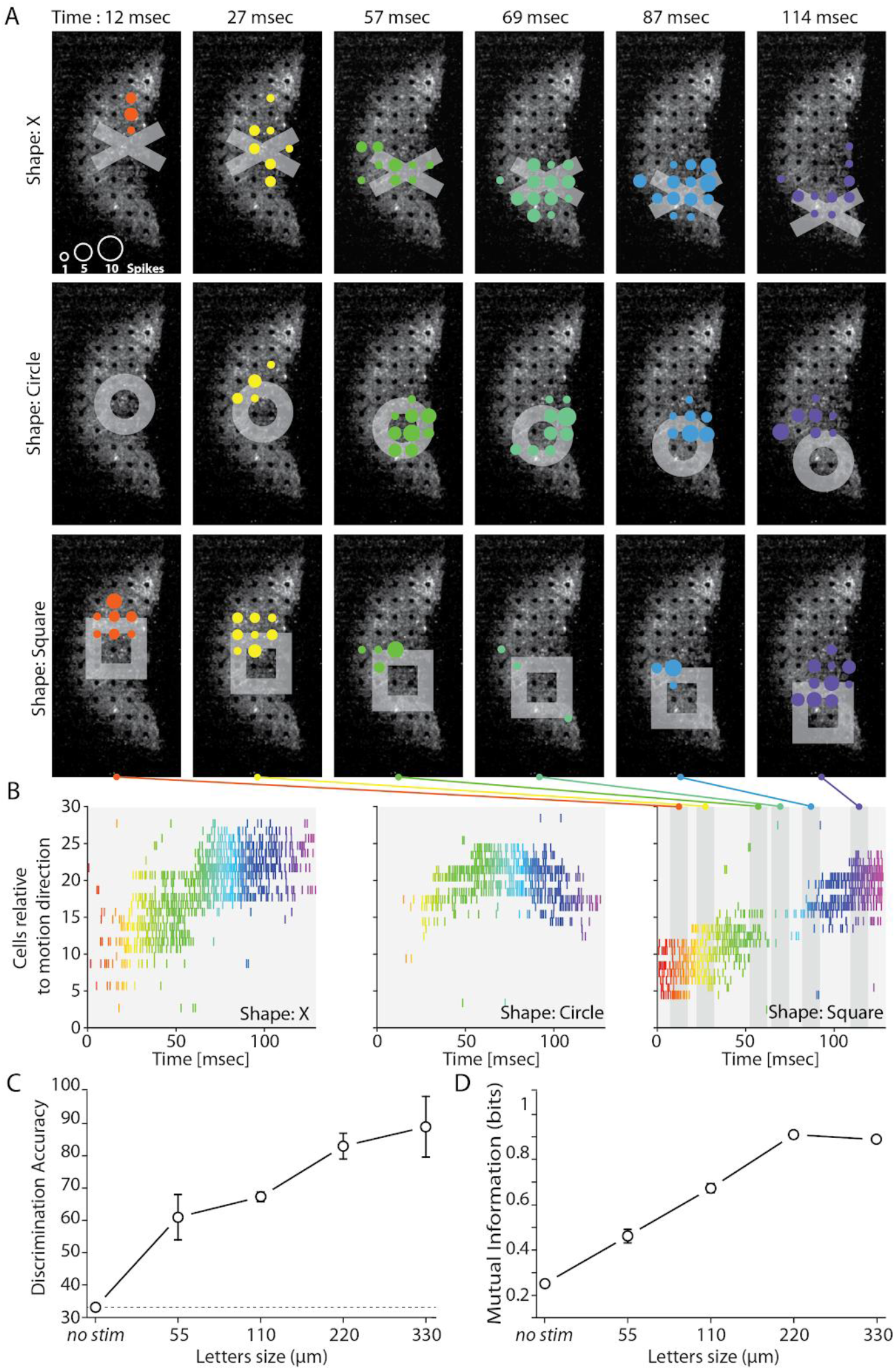
Measured activity can be used to discriminate stimulation with different shapes and predict visual acuity. Three shapes (X, Circle and Square] were moved center-out configuration over the retina. (**A-B**) Response of cells with letters moving from center towards the bottom on the retina. The colormap indicates the time of spike **(A)** and the position of cell on the MEA grid **(B)**. Raster plots show the activity of the cells with time. The three shapes show distinct difference in activity of the population of cells. (**C**) Discriminability (in %) of spatio-temporal spike response features obtained from population spike trains for different sizes. The discriminability improves as the size of letters is increased. No stim recognition was performed using the spontaneous activity pattern just before the start of stimulation. The recognition rate for ‘no stim’ happens to be around 33% i.e. chance level for 3 shapes. (**D**) Normalized mutual information from the confusion matrix obtained from discrimination rates in (C). The information tends to increase with the size of letters saturating at 220 um. Information and high discriminability at 220 um indicates a visual acuity above legal blindness limit.

Due to inherent imperfection in spike sorting and the electrode array sampling only a fraction of transfected cells (~80 cells after spike sorting vs >1000 ChR-tdT cells per retina (Fig. 2)) we are, with no doubt, greatly underestimating the information transmission capabilities of our strategy. Interestingly, we measured for smaller stimuli a steady increase in discrimination accuracy and information (Fig. 7C-D), with access to the complete information transmitted by RGCs we would likely have higher discrimination rate for smaller size symbols.

Despite this limitation, we demonstrate here the ability of our strategy to encode information about speed and direction of stimuli, even for fast moving small stimuli, as well as supporting a discrimination task.

## Discussion

We established in the present study the best efficacy of AAV2.7m8 ChR-tdT construct in primate retinal ganglion cells with respect to the wild type capsid AAV2 and to the non-fused protein ChrimsonR. Furthermore, we defined the therapeutic vector dose at 5 × 10^11^ vg/eye, a dosage that allows for a greater light sensitivity, an expression in more cells and on a wider area. We explored key parameters of optogenetic activation (i.e. light intensity, temporal and spatial modulation) and demonstrated the ability to decode from the recorded cell activity the direction and speed of a moving bar as well as the identity of different stimuli shapes. These data have supported the application to start the ongoing clinical trials on patients affected by retinitis pigmentosa^37^.

### Vector optimization for high functional efficacy

To be successful and reach our therapeutic goals, the viral optogenetic construct must demonstrate high functional impact on a large fraction of the RGCs population. Our results confirm the enhanced transduction efficacy of the AAV2.7m8 variant in the retina with respect to the previous studies using AAV2^8,20^. Surprisingly, in our study we also found a higher efficacy for the fused protein ChR-tdT than for ChR alone. We cannot fully exclude an experimental bias as ChR expression cannot be localized by fluorescence imaging as for ChR-tdT. However, the expected location of the transduced gene at the fovea^20,25^ greatly reduce the chances of missing cluster of expression during the tissue isolation and positioning on the MEA. A more likely explanation resides in differential protein trafficking with td-Tomato working as a trafficking helper, as well as possibly preventing protein aggregation^38^. The result would be an increased membrane targeting of the opsin construct^39^. We demonstrated the greater efficacy for both the mutated AAV capsid, AAV2.7m8, and the fused protein, ChR-tdT, in non-human primates. Due to very high structural similarities between the eye in NHP and humans, we expect that an AAV2.7m8-ChR-tdT intravitreal injection would also effectively transduce the retina in blind patients

### Dose selection

Once the construct is selected, the next critical issue is AAV safety. Previous studies in gene therapy within the eye have used AAV doses of 1.5 × 10^11^ vg in the subretinal space^40^ and up to 1 × 10^11^ vg/eye^41^ or 1.8 × 10^11^ vg/eye^42^ during intravitreal injections. While at these doses no aversive effect was described, other delivery methods use much higher title (up to 10^14^ vg) that can have disruptive effects^43^. In the present study, none of the injected eyes displayed concerning inflammatory responses (fig. S1), with only a few cells in the vitreous and one eye showing vitreal haze (that particular instance was associated with a light hemorrhage during IVT procedure). We observed that 5 × 10^10^ and 5 × 10^11^ vg/eye were very efficient doses to induce ChR-tdT expression with high functional responses in NHPs. The highest dose used here (5 × 10^11^ vg/eye) appears to provide more extensive retinal coverage and higher light sensitivity (Fig. 2). This increased coverage would enlarge the patient’s visual field, which would translate to a ~6 degree in the visual field (211 μm on the primate retina per degree angle,^34^). While this visual field may seem rather limited, it is important to remember that the fovea is the center of high visual acuity. This high visual acuity results from the large density of cone photoreceptors, and the large predominance of midget RGCs, which receive inputs from a single photoreceptor and have very reduced receptive field^44–47^ Using spike propagation speed analysis (fig. S2) and morphology examination, we demonstrated that most of ChR-tdT expressing RGCs are indeed midget RGCs, likely to generate high acuity visual perception. Specific activation of this midget RGC population in the foveal area has the potential to provide patients with a high acuity vision.

### Effective and safe stimulation intensity

We showed here that the stimulation of ChR-tdT, as for most microbial opsin, is effective from 10^15^ photons.cm^-2^.s^-1^. If we consider that only a quarter of the visible light is effective in stimulating any given light sensitive channel, it is hard to find any everyday life situation that would activate it: outdoors during a bright day the effective retinal intensity would be around 10^14^ photons.cm^-2^.s^-1^, and 2×10^12^ photons.cm^-2^.s^-1^ in the office. Thus, we don’t expect the transfected opsin to generate a useful perception when used alone. It is thus necessary for our strategy to include an external photostimulation device which will convert images into tailored patterned photostimulation on the optogenetically engineered retina.

The present study was centered on the red-shifted opsin ChrimsonR with reported peak sensitivity at 575nm^23^, a result we confirmed here using MEA recordings (Fig. 1E). These wavelengths are much safer to use than the highly phototoxic blue wavelength, and as such allow one to safely expose the retina to higher light intensities^29^. For the clinical trial, we decided on a middle ground between the opsin sensitivity and reduced phototoxicity of higher wavelengths and selected a 595 nm LED (Cree XP-E2, Lumitronix) as the light source for the external photostimulation device. For this reason we used 600 nm (+/− 10 nm) light in most of the study, and consider this wavelength for the safety evaluation.

The international commission on non-ionizing radiation protection published limits of ocular exposure for visible and infrared radiaton in 2013^28^, these limits were translated into retina irradiace in Sengupta et al. using previsouly published conversions^48,49^ The resulting threshold was, for continuous exposition, 5.6 × 10^17^ photons.cm^-2^.s^-1^ at 590 nm and 5 × 10^15^ photons.cm^-2^.s^-1^ at 500nm. In 2016, Yan et al. published a review of the 2014 ANSI Z136.1 exposure limits for laser retinal illuminations by ophthalmic instruments (§8.3) and how these limits could be employed in optogenetic systems^21,30^. For a full-field continuous stimulus applied during 8 hours per 48 hours, the maximal permissive retinal peak irradiance is 1.1 × 10^17^ photons.cm^-2^.s^-1^ at 600 nm and 8 × 10^15^ photons.cm^-2^.s^-1^ at 505 nm.

Both sets of proposed exposure limits are higher for the 600nm wavelength used here than for the blue light used for opsins such as Chr2. Importantly, we demonstrate in this paper that light intensities below the 600 nm radiation safety limit can activate the transfected retina very efficiently.

Exposure limits are, by nature, conservative. They concern here a continuous illumination, as the higher risk is the photochemical hazard, which depends on the total illumination received per 48 hours. But our strategy will rely on light patterns stimulation extracted from an event camera reducing moving faces and objects to their outlines^50,51^. The scarcity of stimulation will increase the maximal permissive retinal peak irradiance^21^.

It is also not clear what risk would exist of a photochemical injury in an advanced RP patient’s retina, where the photoreceptors are not functioning and the build-up or potentially toxic retinoids in the RPE would presumably be very minimal.

To further guarantee patient safety during the clinical trial, it is important to initially limit the exposure duration and to closely monitor the state of the retina after each period of stimulation

### Comparison with other studies and constructs

A previous clinical trial in optogenetic therapy for visual restoration had selected the blue-sensitive microbial opsin, Chr2^9^. This first clinical study relied on preclinical studies in mice^52^ and marmosets^53^. In the later study, a single MEA electrode recorded spike trains reaching >300 Hz at 6.6 × 10^16^ photons.cm^-2^.s^-1^. More recently using the human codon-optimized Ca2+-permeable Chr2, (CatCh) – which is 70 times more efficient than Chr2 –, we recorded in macaque foveal RGCs multiunit spiking frequencies in a similar range (~300 Hz at 8 × 10^15^ photons.cm^-2^.s^-154^. These studies rely on opsins sensitive to blue wavelength, and reported results at intensities above safety limits. In the present study, we observed multi-unit spiking frequencies above 700 Hz (Fig. 3A) at 2 × 10^17^ photons.cm^-2^.s^-1^ and peak firing rate above 300Hz for light intensity well above the safety limits (9 × 10^15^ photons.cm^-2^.s^-1^, Fig. 2C).

ChrimsonR is currently the most red-shifted available opsin, ~100 nm from ChR2 and 45 nm from ReaChR peak sensitivities^23^, but future studies may provide even infrared-sensitive opsins as in snakes^55^. Alternatively, mutagenesis can be used to enhance existing opsins properties: ChrimsonR kinetics are the result of directed mutagenesis on Chrimson^23^, and was further modified to drive neuron firing rate to higher spiking frequencies^56^. This kinetics enhancement came at the cost of reduced light sensitivity, and therefore this new variant is not relevant for vision restoration as it would reduce the safe range for stimulation. Considering previous studies aiming toward the development of an optogenetic vision restoration strategy, our results demonstrate the highest level of evoked activity for a 600nm wavelength, with milliseconds temporal resolution at intensity below safety threshold.

### Pattern recognition for visual restoration

Visual restoration with retinal prostheses was classically assessed by stimulating individual electrodes^57,58^, defining object positions and shape^5,58–60^, identifying bar orientation^5,57,59^ and reading letters or word. An issue with the epiretinal prostheses is that single electrode stimulations activated RGCs axons on their way to the optic. This lead to patients reporting perception of an arc instead of a point^61^. In our optogenetic approach, although RGCs axons expressed ChR-tdT, indicated by tdT fluorescence, the applied light stimulation was not able to trigger spikes (Fig. 5B-C). This specificity of activation is a requirement for spatially restricted stimulation; here we show through spot stimulations that responses can be elicited with 50 μm spot diameter even with 10 ms duration (Fig. 5 and fig. S3). While very encouraging, this measure underestimates the actual spatial resolution for two reasons. First, the circular spot used in those experiments were centered on the 10 μm opaque electrodes, reducing photon flux, especially for smaller stimuli (i.e. 25 μm, Fig. 5). Second, due to sampling limitation while recording in MEA, it is likely we did not record the activity of all responsive cells. Therefore, it is somewhat difficult to compare our results to subretinal implants because most of these prosthetic devices activate RGCs indirectly through bipolar cells. However, electrodes have a 70 to 100 μm pitch which can only be activated in a step wise manner as the whole or most of the implant electrode needs to be lit for it to generate the activation current^5,62^ (+ Prevot et al., 2019) whereas optogenetic therapy can activate RGCs with smaller spot size and increasing the size of the stimulation spot increases the number of recruited cell nearly linearly.

Visual acuity is considered normal for a value of 20/20, which correspond to the ability to identify an object of 5 arc minutes, with a critical gap that needs to be resolved of 1 arc minute. The best reported visual acuity with current retinal prostheses was 20/546 when assessed with Landolt C-rings^5^. In our optogenetic strategy, using an approach similar to the Snellen chart we showed correct shape recognition (Fig. 7: 83% recognition for symbols of 220μm with 44 μm edges). Considering that for adult *macaca fascicularis* 1 arc degree in the visual field correspond to a size of 211 μm on the retina^34^, to obtain a 20/20 acuity the size of the gap that need to be resolved is 3.6 μm. In our case the size of gap correctly discriminated (edge of the symbol: 44 μm) represent a 20/249 visual acuity, which is above the legal blindness limit (20/400,^63,64^). This value is in agreement with simulation predictions^65^, and better than any reported acuity with visual prosthetics (20/546,^5^). Our inability to record all cells from the ChR-tdT expressing population clearly undermines this estimation of the visual acuity, which could be even further enhanced. One important perifoveal feature that we did not take into account here is the lateral displacement of RGCs cell body with respect to their receptive field. Indeed to ensure minimal optical aberration for the light hitting the photoreceptor in the most central part of the fovea, other retina layers are displayed centrifugally around the fovea pit. Due to this displacement, the direct stimulation of RGCs should be done through a corrected image of the visual field.

Using reliability of generating spike train, we showed that pulse width modulation might be preferable to continuous illumination (Fig 4). However, if evaluation in patients will define the best stimulation duration, from our data we can infer a reliable activation of RGCs in a 5 to 20 ms range (Fig. 3 & 4). We hypothesize that this ability to evoke high-frequency modulation would help 1) to reduce the total amount of light entering the eye, and 2) to maintain precise control of cell activity. This will result in a more favorable outcome for the strategy.

## Conclusion

The work described here presents an important initial selection process for the genetic content, vector serotype and vector dose in an ambitious “two prongs” vision restoration strategy: a biological component and an external light stimulation device. Our results here, on the biological component, provide all the essential information required for the design of the external light stimulation device necessary for the conversion of the visual scene into a stimulation pattern. Both parts are now in use, combined, in the phase I/II clinical trial that recently started using AAV2.7m8-ChR-tdT^37^

## Materials and Methods

### Animals

Experiments were performed on 21 male and female crab-eating macaques *(Macaca fascicularis)*. The exact number of animals included in the different experiments is listed in table S1. All experiments were done in accordance with the National Institutes of Health Guide for Care and Use of Laboratory Animals. The protocol was approved by the Local Animal Ethics Committees and conducted in accordance with Directive 2010/63/EU of the European Parliament.

### Statistical analysis

All data shown in figures are expressed as mean +/− standard deviation. A *P* value below 0.05 was considered significant. In the test of different genetic constructs, the fraction of active and responsive electrodes was compared between groups using a Chi-square contingency test, followed by Fisher’s exact test (Fig. 1C). When calculating RGC density as a function of eccentricity we compared 5 × 10^09^ and 5 × 10^11^ to 5 × 10^10^ vg/eye for each eccentricity using Tukey’s multiple comparisons test (Fig. 2B). For the percent of responsive electrodes at the different virus concentrations and the two expression times, we performed, for each pair of conditions, a Mann-Whitney non parametric test (Fig. 2, D). The amount of added firing rate for stimulations at different light level for retina treated with three doses of vectors was compared using Tukey’s multiple comparisons test (Fig. 2E). To compare the size of photocurrent in patch clamp and the firing rate in cell attached for the two expression time (Fig. 4B), we performed Mann-Whitney non parametric test. For spot stimulation (Fig. 5) we performed pair-wise t-test over spike rate data for the recorded retinas over different stimulation.

## List of Supplementary Materials

### Supplementary Materials and Methods

**Supplementary Figures**

**Fig S1. RGCs total count in NHP fovea.**

**Fig. S2. Clinical evaluation of inflammation after AAV2.7m8-ChR-tdT treatment.**

**Fig. S3. Transfected RGCs are almost entirely midget cells.**

**Fig. S4. Electrode wise stimulation on a sample retina.**

**Fig. S5. Plane fitting for optical flow calculation.**

**Table S1. Information on all animals included in the study.**

**Table S2. Results of Tukey’s multiple comparisons test for doses responses effect.**

## Acknowledgments

We thank Matthew Chalk for critical proofreading. We thank Guillaume Labernède and Antoine Rizkallah for RGCs manual counts.

## Author contributions

DP-PH-JAS-DD-JD-RB-SP designed the study, AD-MD-DD designed viral vectors, MD produced viral vectors, SB-EB perform IVT injection CMF-JD-CJ-EB performed clinical follow-up on animals, PP-FA-MAK contributed to the in vivo tests, JC controlled all illumination devices, VF prepared the retina according to safety rules, GG-HA-RC-OM designed, performed and analyzed MEA experiments, AC designed, performed and analyzed patch clamp experiments. GG-HA-AC constructed the figures GG-HA-AC-FA-JC-PP-SP-wrote the paper. All authors reviewed the paper.

## Data and materials availability

All data associated with this study are in the paper and/or the Supplementary materials.

