## Supplementary Materials for "Optogenetic therapy: High spatiotemporal resolution and pattern recognition compatible with vision restoration in non-human primates"

#### Supplementary Materials and Methods

##### *AAV production*

ChrimsonR and ChrimsonR-tdTomato were cloned into an AAV backbone plasmid. The constructs all included WPRE and bovine growth hormone polyA. Recombinant AAVs were produced by the plasmid co-transfection method [66], and the resulting lysates were purified via iodixanol gradient ultracentrifugation as previously described. Briefly 40% iodixanol fraction was concentrated and buffer exchanged using Amicon Ultra-15 Centrifugal Filter Units. Vector stocks were then tittered for DNase-resistant vector genomes by real-time PCR relative to a standard [67].

##### *Gene delivery, retina isolation and preservation of the primate retina*

Primates were anesthetized using 10:1 mg/kg of a ketamine/xylazine mix. 100µL of viral vector solution was injected into the vitreous. Following the injection an ophthalmic steroid and antibiotic ointment was applied to the cornea. 2 months (+/- 5 days) or 6 months (+/- 9 days) after AAV injection, primates received a lethal dose of pentobarbital. Eyeballs were removed and placed in sealed bags for transport with $CO_2$ independent medium (ThermoFisher scientific), after puncturation of the eye with a sterile 20-gauge needle. Retina were then isolated and conserved as retinal explants in an incubator for 12 to 36 hours prior to recording. Hemi-foveal retinal fragments were transferred on polycarbonate transwell (Corning) on Neurobasal + B27 medium for conservation in the cell culture incubator.

***Two-photon live Imaging and single-cell electrophysiological recordings***

A custom-made two-photon microscope equipped with a 25x water immersion objective (XLPLN25xWMP, NA: 1.05, Olympus) with a pulsed femto-second laser (InSight™ DeepSee™ - Newport Corporation) was used for imaging ChR-tdT-positive retinal ganglion cells. AAV-treated macaque retinas were imaged in oxygenized (95% O2, 5% CO2) Ames medium (Sigma-Aldrich). For live two-photon imaging, whole-mount retinas were placed in the recording chamber of the microscope (with ganglion cell layer side up), and images and z-stacks were acquired using the excitation laser at a wavelength of 1050 nm. Images were processed offline using ImageJ.

We used an Axon Multiclamp 700B amplifier to perform whole-cell patch-clamp and cell-attached recordings. Patch electrodes were made from borosilicate glass (BF100-50-10, Sutter Instruments) and pulled to 6-9 MΩ. Pipettes were filled with 112.5 mM CsMeSO4, 1 mM Mg SO4, $7.8 \times 10^{-3}$ mM CaCl2, 0.5 mM BAPTA, 10 mM HEPES, 4 mM ATP-Na2, 0.5 mM GTP-Na3, 5 mM lidocaine N-ethyl bromide (QX314-Br) (pH 7.2). We clamped the cells at a potential of -60 mV (the reversal potential of $Cl^-$) in order to isolate excitatory currents. Recordings were also performed in cell-attached configuration, with pipettes filled with Ames medium. Retinas were dark-adapted at least 30 minutes in the recording chamber prior to recordings. Recordings were done in the presence of a selective group III metabotropic glutamate receptor antagonist, L-(+)-2-Amino-4-phosphonobutyric acid (L-AP4, 50 μM, Tocris Bioscience, Bristol, UK).

***MEA***

Multitude Electrode Array (MEA) recordings were obtained from retinal fragments placed on a cellulose membrane preincubated with polylysine (0.1%, Sigma) overnight. Once on a micromanipulator, the retinal piece was gently pressed against a MEA (MEA256 100/30 iR-ITO; Multi-Channel Systems, Reutlingen, Germany), the retinal ganglion cells facing the electrodes. tdTomato fluorescence, when present, was checked prior to recordings with a Nikon Eclipse Ti inverted microscope (Nikon, Dusseldorf, Germany) mounted under the MEA system. The retina was continuously perfused with Ames medium (Sigma-Aldrich, St Louis, MO) bubbled with 95% $O_2$ and 5 % $CO_2$ at 34 °C at a rate of 1–2 ml/minute during experiments. AMPA/kainate glutamate receptor antagonist, 6-cyano-7-nitroquinoxaline-2,3-dione (CNQX, 25 μM, Sigma-Aldrich), NMDA glutamate receptor antagonist, [3H]3-(2-carboxypiperazin-4-yl) propyl-1-phosphonic acid (CPP, 10 μM, Sigma- Aldrich) and a selective group III metabotropic glutamate receptor agonist, L-(+)-2-Amino-4-phosphonobutyric acid (L-AP4, 50 μM, Tocris Bioscience, Bristol, UK) were freshly diluted and bath applied through the perfusion system 10 minutes prior to recordings. Action potentials were detected on the filtered electrode signal (2$^{nd}$ order high pass Butterworth, cut off frequency 200Hz) using a threshold of 4 x SD of the signal. Spike density function, calculated as in [68,69], was averaged over stimuli repetitions and used to measure maximal firing rate in a window corresponding to stimulus duration plus 50 ms. When comparing responses for different vector constructs at different light intensities, we computed for each electrode the added firing rate as the maximal firing rate minus the spontaneous firing rate for this electrode which is calculated as the average firing rate in the 2 seconds prior to stimulation.

##### *Photostimulation*

For single cell electrophysiological recordings, photostimulation was done using a Polychrome V monochromator (Olympus, Hamburg, Germany) set to 600nm (+/- 10 nm) Output light intensities were calibrated to range from $5.8x10^{14}$ to $3.15x10^{17}$ photons.$cm^2$.$s^{-1}$, by using a spectrophotometer (USB2000+, Ocean Optics, Dunedin, FL). A customized videoprojector (Acer DLP P1223, Taiwan) in combination with a chromatic filter (590BP20-nm) was used for randomized oscillating stimulus (Fig. 4J).

For MEA recordings, full field light stimuli were applied with a Polychrome V monochromator (Olympus, Hamburg, Germany) set to 600nm (+/- 10nm), driven by a STG2008 stimulus generator (MCS). Output light intensities were calibrated to range from 1.37 x $10^{14}$ to 6.78 x $10^{16}$ photons.$cm^2$.$sec^{-1}$. For intensity curves we used 2 seconds flashes for 5 Intensities (1.37 x $10^{14}$ , 6.56 x $10^{14}$, 2.34 x $10^{15}$, 8.82 x $10^{15}$, 6.78 x $10^{16}$ photons.$cm^2$.$sec^{-1}$), each repeated 10 times with 5-second inter-stimulus interval. The action spectrum was obtained using a randomized sequence of wavelength (400 to 650 nm, using 10 nm steps) at constant light power, each repeated 10 times with 5-second inter-stimulus interval. High temporal precision and pattern stimulation (moving bar or circular spot) were done using a digital micromirror display (DMD, Vialux, resolution 1024x768) coupled with a LED fluorescence microscope light source (Xcite, Lumen Dynamics) in addition with a chromatic filter (600BP20-nm), and calibrated to deliver a maximum of 2 x $10^{17}$ photons.cm².$sec^{-1}$ on the retina. For stimulation duration assay we applied 17 different durations (10, 20, 30, 40, 50, 100, 200, 300, 400, 500, 1000 and 2000 ms) each repeated 10 times with 5-second inter-stimulus interval.

***Confocal Imaging and quantification***

After MEA experiments, the tissue was recovered and fixed for 30 min at room temperature (PFA 4%), rinsed with PBS and stock at 4°C in sodium azide. 3 half fovea per dosage condition were fixed without prior MEA recordings to prevent possible damage to the retina that would impede cell counting. These retinas were fixed on the same day as the MEA experiments on the same eye and other half of the fovea. Hemi-foveas were then mounted in DAPI containing Vectashield (H-1000, Vector Laboratories) between slides and coverslip (18 x 18 mm, Biosigma) using a 100 µm spacer (Secure-seal space S24735, ThermoFisher Scientific), and then sealed using nail polish. The retinas were imaged on an inverted confocal (Fluoview 1200, Olympus), with a 20X objective (UPLSAPO 20XO, NA: 0.85, Olympus), voxels size comprised between 0.265 and 0.388 µm/pixel in x and y and 1.64 µm/pixel in z. For each hemi-fovea multiple stack were registered and recomposed in an automatic stitch (10% overlap). Using Td-tomato fluorescence we performed blind manual 3D count of the transfected cells in ImageJ (http://imagej.nih.gov/ij) cell counter plugin. The results were then processed using custom made matlab analysis software, allowing *local density* calculation, the *density relative to eccentricity* and the estimation of the *averaged total transfected cells*. *Local density calculation* was performed using binning on cells coordinates, the size of individual bins was determined to be as close as possible to 50µm² while also having a whole number of bin for every stitch. The *density relative to eccentricity* is calculated using Euclidian distances of each counted cell from the fovea center. To estimate the fovea center position we used the center of a disk drawn to best describe the fovea shape. Our retina explants were cut in order to have hemi-foveas on two explants, but the angle produced by the cutting process was not 180 degree. For each explant we estimated the angle covered by the retina explants using the fovea center we calculated, and we used that number to estimate the total density of ChR-tdT expressing RGCs relative to eccentricity.

The *averaged total transfected cells* estimate is the total number of counted ChR-tdT positive cells for an explants divided by the angle of fovea covered in the explants and then multiplied by 360 degrees.

***Descriptive statistics of population response on multielectrode array***

When analyzing response pattern for stimuli of increasing duration (1 ms to 2000 m, Fig.3) we compute the *time to first spike* and *Fano factor*. Time to first spike is determined as the time span between initiations of stimulation and the next spike on the electrode. For each electrode that value is averaged over all repetitions for a specific duration. We then build a box-plot for each duration containing all electrodes in all experiments. The Fano Factor, *F*, was calculated for each electrode as:

$$F = \frac{< \delta N^2 >}{< N >}$$

Where $< \delta N^2 >$ the average variance of spike is divided by $< N >$, the average spike count.

***Letter recognition using population response***

Three different shapes (X, circle and square) were presented over the retina moving in center-out configuration in eight directions. The shapes were created using the Snellen chart ratios such that the ratio of width of the individual edges of the shape to the total size was 1:5. The widths were set from 11 um to 66 um creating shapes of size 55um to 330um. Each condition was presented over 50 trials. The response of the population of cells for each representation could be represented as a set of $K$ spikes $S = \{S_1, S_2, S_k \dots S_K\}$. Each spike $S_k$ is a 2-dimensional word $\{i, t_k\}$, representing a spike at cell $i$ at time $t_k$.

The spike train $S$ can be considered as a population signature representing each presented stimuli. These spike patterns $S$ were used to perform pattern recognition similar to an algorithm [70] developed for event-trains from neuromorphic sensors. The inter-cell spike intervals were used to create temporal features. For each set $S$, a feature vector $F \in R^N$ ($N$ = Number of cells) was created by averaging the temporal feature vectors $f$ for each spike $S_k$ (Eq 1).

$$f_{k,i} = e^{-\frac{|t_k - t_i|}{\tau}} \qquad (1)$$

$$F(i) = \frac{1}{K}\sum_{k=1}^{K} f_{k,i} \qquad (2)$$

where $t_k$ is the spike time for $S_k$ and $t_i$ is the last spike at every other cell $i$ and $\tau$ is the decay constant that dictates the effect of temporal history of the last spike of the cells and was set to 5 ms in this case. Thus, each trial $T$ of a shape is represented by the vector $F$. Finally, a k-means based linear discriminator, trained over half of the trials was used to discriminate the remaining half of the trials into one of the three shapes. The performance of the feature and k-means was quantified as the mutual information between the presented and predicted shape stimuli, $I(L; L^p)$,

(Eq 3) [71] where $P(L,L^p)$ represents joint probabilities for presented shape $L$ that was predicted by the decoder as $L^p$.

$$I(L;L^p) = \sum_{L,L^p} P(L,L^p)\, log_2 \frac{P(L,L^p)}{P(L)P(L^p)} \qquad (3)$$

This measure is more useful than the average discrimination accuracy as it also incorporates cross stimulus errors. In effect, it provides quantification of both the how close are the features within the same class cluster and how distance they are from the features of all other classes. The above computation was performed using the information breakdown toolbox in matlab [72].

***Moving bar stimulation and direction estimation***

The retina was stimulated with moving bars at two speeds (2.2 mm/sec, 4.4 mm/sec) and four orientations (0, 45, 90, 135$^o$) (Fig. 8). Each trial consisted of the motion of the bar across the whole retina. A total of 200 trials were recorded for each condition. Each trial creates a 3-dimensional pattern of spikes where each spike can be represented as $R^3 = \{x, y, t\}$ where $[x, y]$ represent the index of the MEA electrode location and $t$ is the time of spike from start of bar motion (fig. S5). A simple least square estimate method was used to fit a plane (Eq 4). The partial and composite slopes from the coefficients of the plane provide both the speed and orientation of the bar if the temporal precision of the spikes generated is high enough. That is the cells can respond at a faster rate than the time it takes for the bar to move from one cell to the next.

$$Ax + By + Ct + D = 0 \quad (4)$$

$$\frac{\delta x}{\delta t} \equiv \frac{-C}{A}, \quad \frac{\delta y}{\delta t} \equiv \frac{-C}{B} \; and \quad \frac{\delta x}{\delta y} \equiv \frac{-B}{A} \quad (5)$$

.

### Supplementary Figures

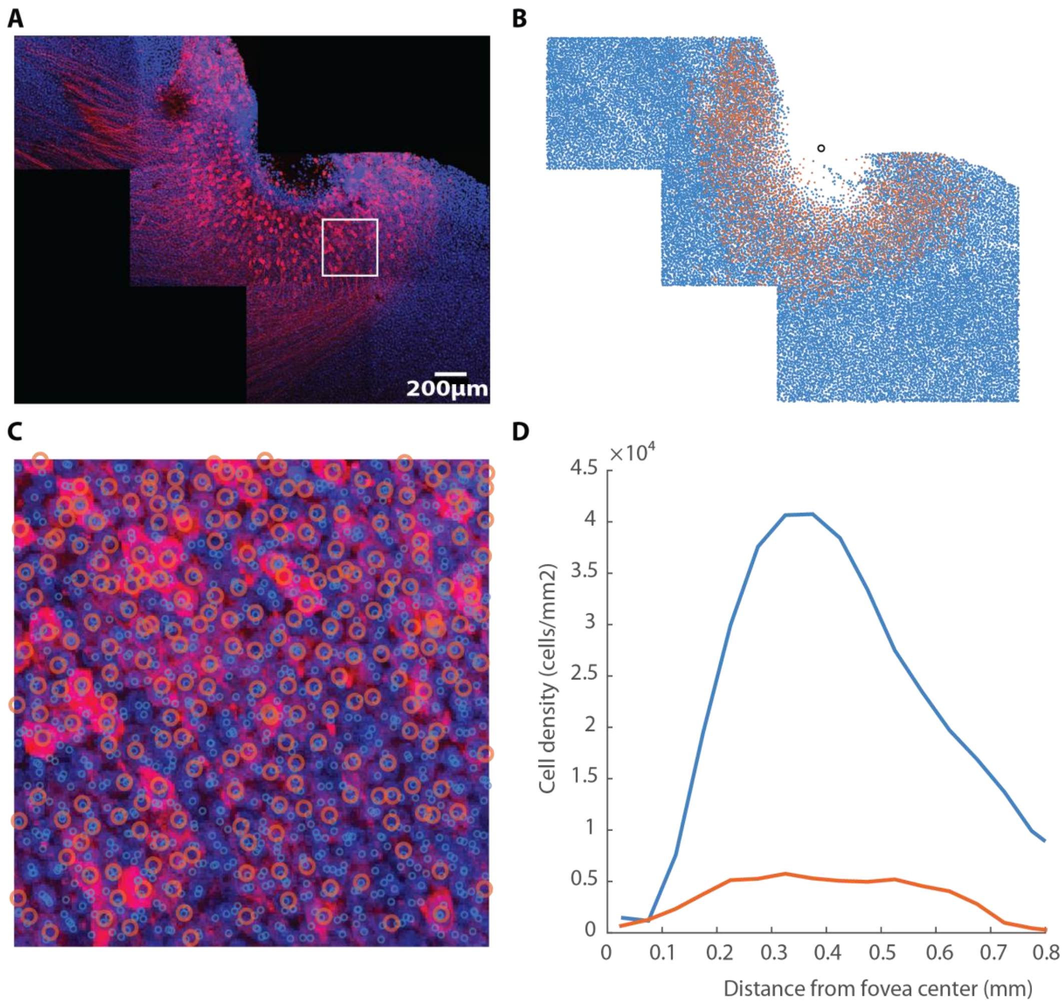


**Fig S1. RGCs total count in NHP fovea.**

**(A)** Projection of stitched confocal stacks of a DAPI stained ChR-tdT expressing fovea. The assembled stitch was edited to remove nuclei below the RGC layer. **(B)** Overlay of manually counted ChR-tdT expressing RGCs and automated nuclei detection using imaris (see supplementary methods) for the complete stitch. The black circle represents the estimated fovea center. **(C)** Close up of the white square shown in (A), open orange circles depict ChR-tdT positive RGCs, blue circles are nuclei labeled by the DAPI staining. (D) Total cell density in the RGC layer relative to retina eccentricity, obtained from DAPI nuclei counts (blue line). Note an apex in density around 40 000 cells/mm² at 0.4mm eccentricity. Density of ChR-tdT positive cells relative to retina eccentricity is provided for comparison (red line), This particular set of images was obtained from a medium dose (5 x $10^{10}$ vg/eye) treated retina and show a maximal density ~5000 cells/mm² (~12.5% of total cells).

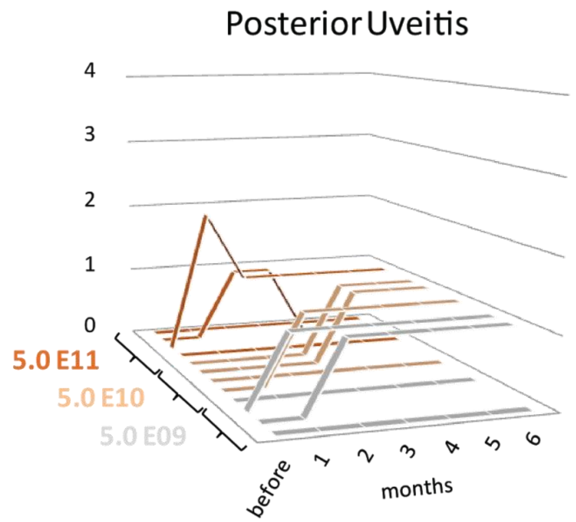


| Grade | Clinical findings |
|---|---|
| 0 | No vitreous cells |
| 1 | Vitreous cells with clear view of posterior fundus |
| 2 | Vitreous cells with poor view of details, and view of the optic nerve and retinal vessel |
| 3 | Vitreous cells with view of optic nerve |
| 4 | Vitreous cells with no view of the posterior segment |

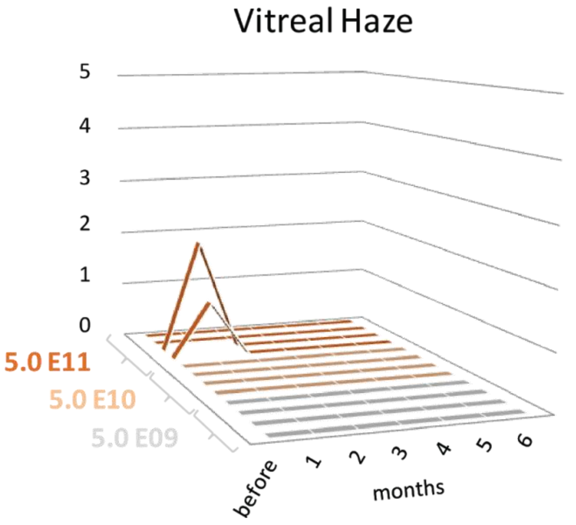


| Grade | Clinical findings |
|---|---|
| 0 | None |
| 1 | Posterior pole clearly visible |
| 2 | Posterior pole details slightly hazy |
| 3 | Posterior pole details very hazy |
| 4 | Posterior pole details barely visible |
| 5 | Fundal details not visible |

**Fig. S2. Clinical evaluation of inflammation after AAV2.7m8-ChR-tdT treatment.**

(***top***) examination for posterior uveitis following the grades described in the table on the right. Note that only one animal (for the high dose of vector) rise to grade 2, and only for the first month. (***bottom***) Report of vitreal haze for the injected animals, only two animals presented signs of haze. The haziest fundus is of the same animal that presented a grade 2 uveitis measurement. The haze dissipated for both animals at 2 months post-injections. These two observations indicate a mild reaction due to the intra-vitreous injection (this particular animal presented a small hemorrhage when the sclera was pierced). The expression of an ectopic transgene usually reaches full potency at 2 months of expression, and does not seem to induce any delayed inflammation.

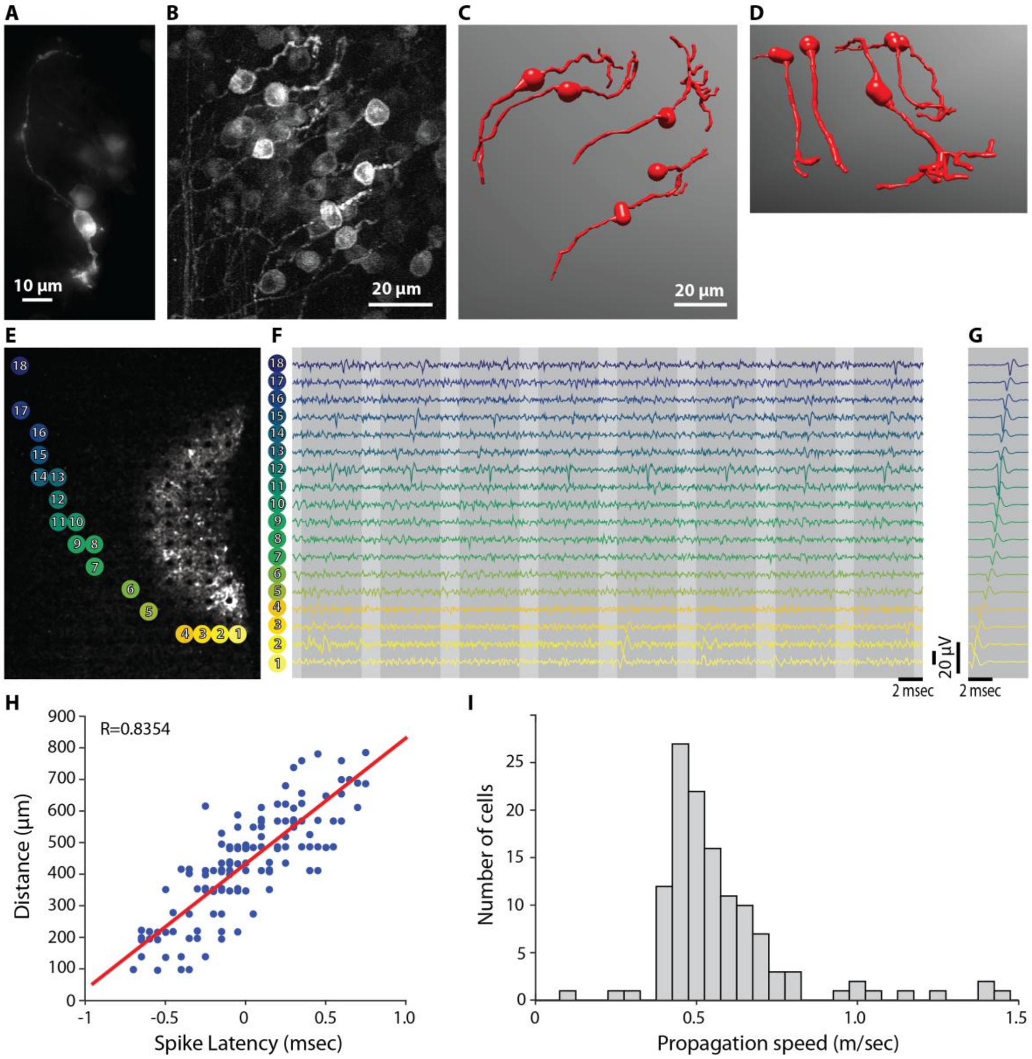


**Fig. S3. Transfected RGCs are almost entirely midget cells.**

(**A**) Epifluoresecence image of a ChR-tdT expressing RGCs close to the fovea, its morphology is typical of midget ganglion: a small soma (circle) with a single dendrite (arrow) and little to no terminal ramification. The axon can be seen on the opposite side of the cell (arrowhead). (**B**) Projection of a 2-photon stack of images displaying a cluster of RGCs with strong ChR-tdT expression. (**C-D**) Bird eye view (C) and side view (D) of a 3D reconstruction of complete RGCs present in the images. 4 of the cells had very restricted terminal dendritic arborization (5-10 μm), while one cell had a larger arborization (~40 μm), indicative of midget ganglion cells and parasol ganglion cells, respectively. Note that the soma of the putative parasol cell is deeper in the ganglion cell layer. (**E-I**) Measurement of spike propagation speed in responsive RGCs expressing ChR-tdT. (**E**) image of the hemi-fovea on the MEA overlaid with the position of the electrodes displaying the responses of one RGCs, numbered 1 to 18, 1 closest to the soma, 18 furthest away. (**F-G**) Traces showing responses in 8 trials (F, dark grey), or the average response (G) to circular stimulation on electrodes 1. On the average response, the progressive increase in spike latency can be clearly seen. (**H**) Linear regression confirms positive correlation between electrodes distance from the site of spike initiation (soma) and the recorded latency at that electrode. The slope of the linear regression was used to estimate the action potential speed propagation for that ganglion cell. (**I**) Histogram for the propagation speed in our population, most of the cells have a propagation speed around 0.5 m/s , confirming they are indeed midget ganglion cells *(32)*. Few cells show greater than 1 m/s propagation speed, those are likely parasol cells.

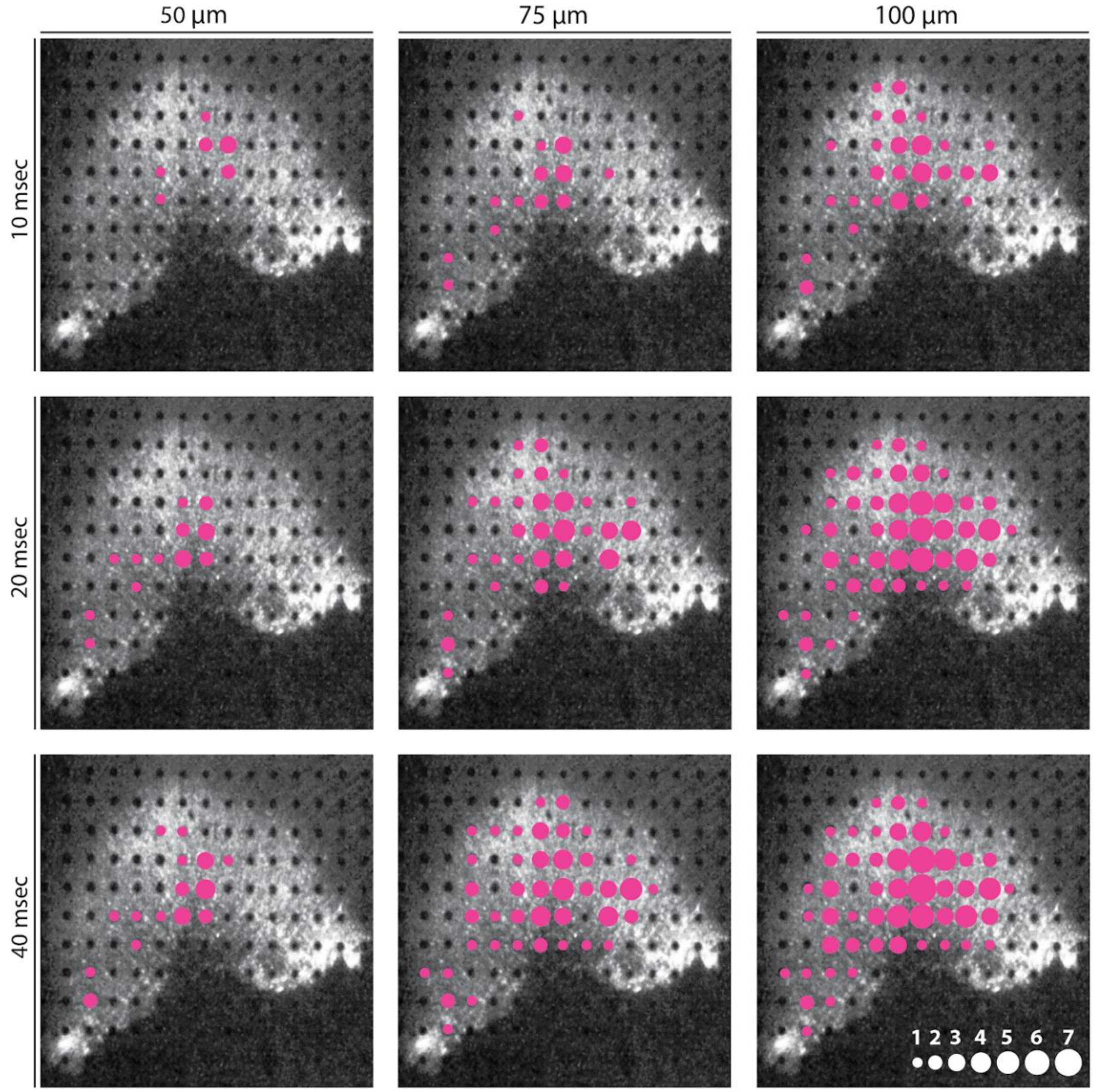


**Fig. S4. Electrode wise stimulation on a sample retina.**

Circular shaped spots of light were presented on the electrode sites with varying duration (10-20-40 ms) and size (50-75-100 µm). Each line of images is a different duration of stimulation, each column a size of stimulation. The number of recruited cell per electrode stimulated is represented as magenta disk of increasing size (see scale in the bottom right corner).

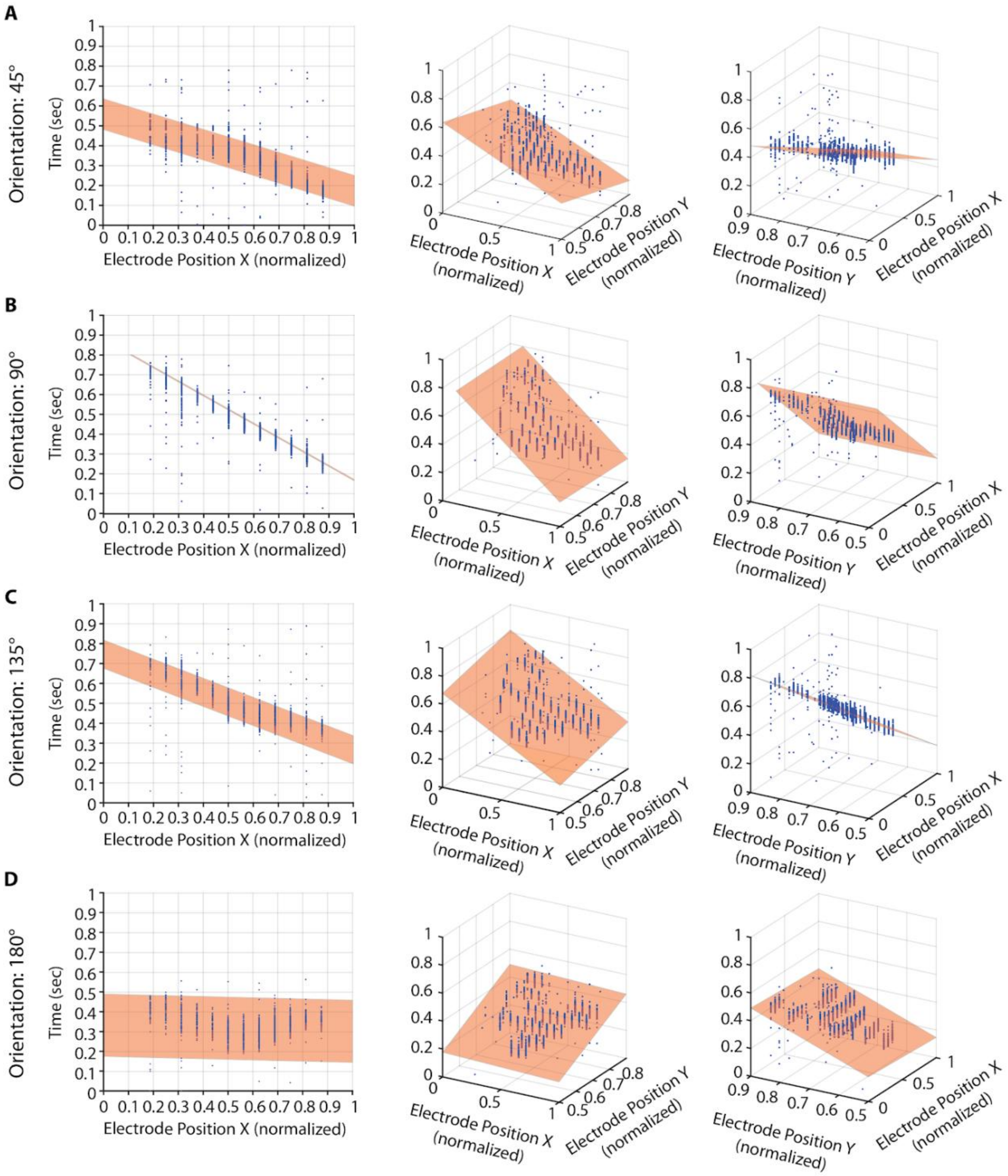


**Fig. S5. Plane fitting for optical flow calculation.**

Four different direction of movement shown here (**A**: 45°, **B**: 90°, **C**: 135°, **D**: 180°). Cell normalized time for peak firing rate is indicated relative to the position of the main electrode recording that cell (Blue dots). The 2D plane fitting the peak activity timing is displayed in orange. Three orientations for the same plot is shown here to better visualize change on the fitted plane (0° rotation on the left, -30° rotation in the middle, and 60° rotation on the right).

| NHP number | Sex | Age (years) | Right eye | | Left eye | | Expression duration (days) |
|---|---|---|---|---|---|---|---|
| | | | Viral genome/ eye | Construct | Viral genome/ eye | Virus | |
| CA662 | M | 3 | 5.00E+11 | AAV2-7m8-ChrimsonR | 5.00E+11 | AAV2-7m8-ChrimsonR-tdTomato | 55 |
| CA680 | M | 4 | 5.00E+11 | AAV2-7m8-ChrimsonR | 5.00E+11 | AAV2-7m8-ChrimsonR-tdTomato | 57 |
| BX914 | M | 4 | 5.00E+11 | AAV2-7m8-ChrimsonR | 5.00E+11 | AAV2-7m8-ChrimsonR-tdTomato | 62 |
| BX430 | M | 4 | 5.00E+11 | AAV2-7m8-ChrimsonR | 5.00E+11 | AAV2-7m8-ChrimsonR-tdTomato | 63 |
| CB206 | F | 7 | 5.00E+11 | AAV2-ChrimsonR | 5.00E+11 | AAV2-ChrimsonR-tdTomato | 55 |
| CA295 | F | 13 | 5.00E+11 | AAV2-ChrimsonR | 5.00E+11 | AAV2-ChrimsonR-tdTomato | 56 |
| CB737 | M | 3 | 5.00E+11 | AAV2-ChrimsonR | 5.00E+11 | AAV2-ChrimsonR-tdTomato | 62 |
| P2 | M | 3 | 5.00E+11 | AAV2-ChrimsonR | 5.00E+11 | AAV2-ChrimsonR-tdTomato | 64 |
| BA586I | M | 4 | 5.00E+10 | AAV2-7m8-ChrimsonR-tdTomato | 5.00E+11 | AAV2-7m8-ChrimsonR-tdTomato | 182 |
| BB161I | M | 5 | 5.00E+09 | AAV2-7m8-ChrimsonR-tdTomato | 5.00E+10 | AAV2-7m8-ChrimsonR-tdTomato | 184 |
| BB478E | M | 4 | 5.00E+09 | AAV2-7m8-ChrimsonR-tdTomato | 5.00E+10 | AAV2-7m8-ChrimsonR-tdTomato | 182 |
| BA901H | M | 4 | 5.00E+11 | AAV2-7m8-ChrimsonR-tdTomato | 5.00E+10 | AAV2-7m8-ChrimsonR-tdTomato | 184 |
| BB222I | M | 4 | 5.00E+09 | AAV2-7m8-ChrimsonR-tdTomato | 5.00E+11 | AAV2-7m8-ChrimsonR-tdTomato | 190 |
| BC110B | M | 5 | 5.00E+11 | AAV2-7m8-ChrimsonR-tdTomato | 5.00E+09 | AAV2-7m8-ChrimsonR-tdTomato | 191 |
| 1107044 | F | 5 | 5.00E+11 | AAV2-7m8-ChrimsonR-tdTomato | 5.00E+11 | AAV2-7m8-ChrimsonR-tdTomato | 177 |
| 1107086 | F | 5 | 5.00E+11 | AAV2-7m8-ChrimsonR-tdTomato | 5.00E+11 | AAV2-7m8-ChrimsonR-tdTomato | 175 |
| 1106077 | M | 5 | 5.00E+11 | AAV2-7m8-ChrimsonR-tdTomato | 5.00E+11 | AAV2-7m8-ChrimsonR-tdTomato | 161 |
| BB325G | M | 4 | 5.00E+11 | AAV2-7m8-ChrimsonR-tdTomato | 5.00E+11 | AAV2-7m8-ChrimsonR-tdTomato | 163 |

**Table S1. Information on all animals included in the study**.

| | | Intensity 4 (8.82 x $10^{15}$ photons.cm$^{-2}$.s$^{-1}$) | | | | Intensity 5 (6.78 x $10^{16}$ photons.cm$^{-2}$.s$^{-1}$) | | | |
|---|---|---|---|---|---|---|---|---|---|
| Dose | Tukey's multiple comparisons test | Mean Diff | 95.00% CI of diff | Summary | Adjusted P Value | Mean Diff2 | 95.00% CI of diff3 | Summary4 | Adjusted P Value5 |
| 5 x $10^9$ vg/eye | (5E09).BB22i_OD vs. (5E10).BA586i_OD | -2.599 | -57.12 to 51.93 | ns | >0.9999 | -36.39 | -90.91 to 18.14 | ns | 0.563 |
| | (5E09).BB22i_OD vs. (5E10).BB478_OG1 | -0.1387 | -56.53 to 56.26 | ns | >0.9999 | -3.324 | -59.72 to 53.07 | ns | >0.9999 |
| | (5E09).BB22i_OD vs. (5E10).BB161i_OG | -8.784 | -61.72 to 44.15 | ns | >0.9999 | -125 | -178 to -72.1 | **** | <0.0001 |
| | (5E09).BB22i_OD vs. (5E11).BA901h_OD | -20.19 | -73.67 to 33.28 | ns | 0.9862 | -215.3 | -268.7 to -161.8 | **** | <0.0001 |
| | (5E09).BB22i_OD vs. (5E11).BB222i_OG | -19.05 | -72.3 to 34.2 | ns | 0.9911 | -118.3 | -171.6 to -65.1 | **** | <0.0001 |
| | (5E09).BB22i_OD vs. (5E11).1106077_OD | -74.26 | -128.6 to -19.88 | *** | 0.0005 | -230.9 | -285.3 to -176.6 | **** | <0.0001 |
| | (5E09).BB22i_OD vs. (5E11).1107086_OD | -116.9 | -171.3 to -62.43 | **** | <0.0001 | -254.4 | -308.8 to -199.9 | **** | <0.0001 |
| | (5E09).BB22i_OD vs. (5E11).1107086_OG | -10.1 | -75.06 to 54.86 | ns | >0.9999 | -42.49 | -107.4 to 22.48 | ns | 0.595 |
| | (5E09).BB22i_OD vs. (5E11).1106077_OG | -4.953 | -64.07 to 54.16 | ns | >0.9999 | -63.51 | -122.6 to -4.388 | * | 0.0228 |
| | (5E09).BB22i_OD vs. (5E11).BB325g_OD | -73.66 | -134 to -13.36 | ** | 0.0038 | -202.7 | -263 to -142.4 | **** | <0.0001 |
| | (5E09).BB22i_OD vs. (5E11).BB325g_OG | -2.548 | -71.26 to 66.16 | ns | >0.9999 | -42.59 | -111.3 to 26.12 | ns | 0.675 |
| Dose | Tukey's multiple comparisons test | Mean Diff | 95.00% CI of diff | Summary | Adjusted P Value | Mean Diff2 | 95.00% CI of diff3 | Summary4 | Adjusted P Value5 |
| 5 x $10^{10}$ vg/eye | (5E10).BA586i_OD vs. (5E10).BB478_OG1 | 2.46 | -29.94 to 34.86 | ns | >0.9999 | 33.06 | 0.6603 to 65.46 | * | 0.0406 |
| | (5E10).BA586i_OD vs. (5E10).BB161i_OG | -6.186 | -32.11 to 19.74 | ns | 0.9998 | -88.65 | -114.6 to -62.73 | **** | <0.0001 |
| | (5E10).BA586i_OD vs. (5E11).BA901h_OD | -17.6 | -44.59 to 9.402 | ns | 0.6003 | -178.9 | -205.9 to -151.9 | **** | <0.0001 |
| | (5E10).BA586i_OD vs. (5E11).BB222i_OG | -16.45 | -43 to 10.09 | ns | 0.6754 | -81.96 | -108.5 to -55.41 | **** | <0.0001 |
| | (5E10).BA586i_OD vs. (5E11).1106077_OD | -71.66 | -100.4 to -42.91 | **** | <0.0001 | -194.5 | -223.3 to -165.8 | **** | <0.0001 |
| | (5E10).BA586i_OD vs. (5E11).1107086_OD | -114.3 | -143.2 to -85.4 | **** | <0.0001 | -218 | -246.9 to -189.1 | **** | <0.0001 |
| | (5E10).BA586i_OD vs. (5E11).1107086_OG | -7.5 | -53.21 to 38.21 | ns | >0.9999 | -6.1 | -51.81 to 39.61 | ns | >0.9999 |
| | (5E10).BA586i_OD vs. (5E11).1106077_OG | -2.354 | -39.29 to 34.58 | ns | >0.9999 | -27.12 | -64.05 to 9.816 | ns | 0.4056 |
| | (5E10).BA586i_OD vs. (5E11).BB325g_OD | -71.06 | -109.9 to -32.26 | **** | <0.0001 | -166.3 | -205.1 to -127.5 | **** | <0.0001 |
| | (5E10).BA586i_OD vs. (5E11).BB325g_OG | 0.05052 | -50.84 to 50.94 | ns | >0.9999 | -6.206 | -57.1 to 44.69 | ns | >0.9999 |
| | (5E10).BB478_OG1 vs. (5E10).BB161i_OG | -8.646 | -38.3 to 21.01 | ns | 0.9985 | -121.7 | -151.4 to -92.06 | **** | <0.0001 |
| | (5E10).BB478_OG1 vs. (5E11).BA901h_OD | -20.06 | -50.66 to 10.54 | ns | 0.5916 | -211.9 | -242.5 to -181.3 | **** | <0.0001 |
| | (5E10).BB478_OG1 vs. (5E11).BB222i_OG | -18.91 | -49.12 to 11.29 | ns | 0.6606 | -115 | -145.2 to -84.82 | **** | <0.0001 |
| | (5E10).BB478_OG1 vs. (5E11).1106077_OD | -74.12 | -106.3 to -41.96 | **** | <0.0001 | -227.6 | -259.8 to -195.5 | **** | <0.0001 |
| | (5E10).BB478_OG1 vs. (5E11).1107086_OD | -116.7 | -149 to -84.47 | **** | <0.0001 | -251 | -283.3 to -218.8 | **** | <0.0001 |
| | (5E10).BB478_OG1 vs. (5E11).1107086_OG | -9.96 | -57.89 to 37.97 | ns | >0.9999 | -39.16 | -87.09 to 8.767 | ns | 0.2408 |
| | (5E10).BB478_OG1 vs. (5E11).1106077_OG | -4.814 | -44.46 to 34.83 | ns | >0.9999 | -60.18 | -99.83 to -20.54 | **** | <0.0001 |
| | (5E10).BB478_OG1 vs. (5E11).BB325g_OD | -73.52 | -114.9 to -32.13 | **** | <0.0001 | -199.4 | -240.8 to -158 | **** | <0.0001 |
| | (5E10).BB478_OG1 vs. (5E11).BB325g_OG | -2.409 | -55.3 to 50.48 | ns | >0.9999 | -39.27 | -92.16 to 13.63 | ns | 0.3875 |
| | (5E10).BB161i_OG vs. (5E11).BA901h_OD | -11.41 | -35.04 to 12.22 | ns | 0.917 | -90.22 | -113.9 to -66.59 | **** | <0.0001 |
| | (5E10).BB161i_OG vs. (5E11).BB222i_OG | -10.27 | -33.38 to 12.85 | ns | 0.9529 | 6.691 | -16.42 to 29.81 | ns | 0.9986 |
| | (5E10).BB161i_OG vs. (5E11).1106077_OD | -65.47 | -91.09 to -39.86 | **** | <0.0001 | -105.9 | -131.5 to -80.28 | **** | <0.0001 |
| | (5E10).BB161i_OG vs. (5E11).1107086_OD | -108.1 | -133.9 to -82.33 | **** | <0.0001 | -129.3 | -155.1 to -103.6 | **** | <0.0001 |
| | (5E10).BB161i_OG vs. (5E11).1107086_OG | -1.314 | -45.12 to 42.49 | ns | >0.9999 | 82.55 | 38.74 to 126.4 | **** | <0.0001 |
| | (5E10).BB161i_OG vs. (5E11).1106077_OG | 3.831 | -30.72 to 38.38 | ns | >0.9999 | 61.53 | 26.98 to 96.08 | **** | <0.0001 |
| | (5E10).BB161i_OG vs. (5E11).BB325g_OD | -64.88 | -101.4 to -28.34 | **** | <0.0001 | -77.7 | -114.2 to -41.15 | **** | <0.0001 |
| | (5E10).BB161i_OG vs. (5E11).BB325g_OG | 6.236 | -42.95 to 55.43 | ns | >0.9999 | 82.45 | 33.25 to 131.6 | **** | <0.0001 |
| Dose | Tukey's multiple comparisons test | Mean Diff | 95.00% CI of diff | Summary | Adjusted P Value | Mean Diff2 | 95.00% CI of diff3 | Summary4 | Adjusted P Value5 |
| 5 x $10^{11}$ vg/eye | (5E11).BA901h_OD vs. (5E11).BB222i_OG | 1.143 | -23.17 to 25.46 | ns | >0.9999 | 96.91 | 72.6 to 121.2 | **** | <0.0001 |
| | (5E11).BA901h_OD vs. (5E11).1106077_OD | -54.06 | -80.77 to -27.36 | **** | <0.0001 | -15.67 | -42.38 to 11.03 | ns | 0.7475 |
| | (5E11).BA901h_OD vs. (5E11).1107086_OD | -96.69 | -123.5 to -69.84 | **** | <0.0001 | -39.11 | -65.95 to -12.26 | *** | 0.0001 |
| | (5E11).BA901h_OD vs. (5E11).1107086_OG | 10.1 | -34.36 to 54.55 | ns | 0.9999 | 172.8 | 128.3 to 217.2 | **** | <0.0001 |
| | (5E11).BA901h_OD vs. (5E11).1106077_OG | 15.24 | -20.12 to 50.61 | ns | 0.962 | 151.8 | 116.4 to 187.1 | **** | <0.0001 |
| | (5E11).BA901h_OD vs. (5E11).BB325g_OD | -53.47 | -90.78 to -16.16 | *** | 0.0002 | 12.53 | -24.78 to 49.84 | ns | 0.9948 |
| | (5E11).BA901h_OD vs. (5E11).BB325g_OG | 17.65 | -32.12 to 67.41 | ns | 0.9918 | 172.7 | 122.9 to 222.4 | **** | <0.0001 |
| | (5E11).BB222i_OG vs. (5E11).1106077_OD | -55.21 | -81.46 to -28.96 | **** | <0.0001 | -112.6 | -138.8 to -86.34 | **** | <0.0001 |
| | (5E11).BB222i_OG vs. (5E11).1107086_OD | -97.83 | -124.2 to -71.44 | **** | <0.0001 | -136 | -162.4 to -109.6 | **** | <0.0001 |
| | (5E11).BB222i_OG vs. (5E11).1107086_OG | 8.952 | -35.23 to 53.13 | ns | >0.9999 | 75.86 | 31.68 to 120 | **** | <0.0001 |
| | (5E11).BB222i_OG vs. (5E11).1106077_OG | 14.1 | -20.92 to 49.12 | ns | 0.9772 | 54.84 | 19.82 to 89.86 | **** | <0.0001 |
| | (5E11).BB222i_OG vs. (5E11).BB325g_OD | -54.61 | -91.6 to -17.63 | **** | <0.0001 | -84.39 | -121.4 to -47.4 | **** | <0.0001 |
| | (5E11).BB222i_OG vs. (5E11).BB325g_OG | 16.5 | -33.02 to 66.03 | ns | 0.9952 | 75.75 | 26.23 to 125.3 | **** | <0.0001 |
| | (5E11).1106077_OD vs. (5E11).1107086_OD | -42.62 | -71.23 to -14.01 | **** | <0.0001 | -23.43 | -52.04 to 5.177 | ns | 0.2375 |
| | (5E11).1106077_OD vs. (5E11).1107086_OG | 64.16 | 18.62 to 109.7 | *** | 0.0003 | 188.4 | 142.9 to 234 | **** | <0.0001 |
| | (5E11).1106077_OD vs. (5E11).1106077_OG | 69.31 | 32.58 to 106 | **** | <0.0001 | 167.4 | 130.7 to 204.1 | **** | <0.0001 |
| | (5E11).1106077_OD vs. (5E11).BB325g_OD | 0.5955 | -38 to 39.2 | ns | >0.9999 | 28.2 | -10.4 to 66.8 | ns | 0.4138 |
| | (5E11).1106077_OD vs. (5E11).BB325g_OG | 71.71 | 20.97 to 122.4 | *** | 0.0002 | 188.3 | 137.6 to 239.1 | **** | <0.0001 |
| | (5E11).1107086_OD vs. (5E11).1107086_OG | 106.8 | 61.16 to 152.4 | **** | <0.0001 | 211.9 | 166.3 to 257.5 | **** | <0.0001 |
| | (5E11).1107086_OD vs. (5E11).1106077_OG | 111.9 | 75.1 to 148.8 | **** | <0.0001 | 190.9 | 154 to 227.7 | **** | <0.0001 |
| | (5E11).1107086_OD vs. (5E11).BB325g_OD | 43.22 | 4.519 to 81.92 | * | 0.014 | 51.63 | 12.93 to 90.33 | *** | 0.0008 |
| | (5E11).1107086_OD vs. (5E11).BB325g_OG | 114.3 | 63.52 to 165.1 | **** | <0.0001 | 211.8 | 161 to 262.6 | **** | <0.0001 |
| | (5E11).1107086_OG vs. (5E11).1106077_OG | 5.146 | -45.96 to 56.25 | ns | >0.9999 | -21.02 | -72.12 to 30.08 | ns | 0.9731 |
| | (5E11).1107086_OG vs. (5E11).BB325g_OD | -63.56 | -116 to -11.1 | ** | 0.0043 | -160.2 | -212.7 to -107.8 | **** | <0.0001 |
| | (5E11).1107086_OG vs. (5E11).BB325g_OG | 7.551 | -54.4 to 69.5 | ns | >0.9999 | -0.1058 | -62.05 to 61.84 | ns | >0.9999 |
| | (5E11).1106077_OG vs. (5E11).BB325g_OD | -68.71 | -113.7 to -23.68 | **** | <0.0001 | -139.2 | -184.3 to -94.2 | **** | <0.0001 |
| | (5E11).1106077_OG vs. (5E11).BB325g_OG | 2.405 | -53.38 to 58.19 | ns | >0.9999 | 20.91 | -34.87 to 76.7 | ns | 0.987 |
| | (5E11).BB325g_OD vs. (5E11).BB325g_OG | 71.11 | 14.08 to 128.2 | ** | 0.0027 | 160.1 | 103.1 to 217.2 | **** | <0.0001 |

**Table S2. Results of Tukey's multiple comparisons test for doses responses effect.** Only the results for the two highest light intensities are given, no significant differences for the 3 others light intensities. Out of the 4 medium dose retina, one showed no light responses, two retinas had significantly smaller responses than 5 and 6 (out of 8) high dose retina, at the highest light intensity. Finally, only a single medium dose retina ((5E10).BB161i_OG) has a significantly higher response than 3 of the 8 high dose retinas for the highest light intensity only, demonstrated as a positive mean difference (bold red), it also has a significantly lower response than 4 of the high dose retinas.
